# A genomic analysis and transcriptomic atlas of gene expression in *Psoroptes ovis* reveals feeding- and stage-specific patterns of allergen expression

**DOI:** 10.1101/578120

**Authors:** Stewart TG Burgess, Edward J Marr, Kathryn Bartley, Francesca G Nunn, Rachel E Down, Robert J Weaver, Jessica C Prickett, Jackie Dunn, Stephane Rombauts, Thomas Van Leeuwen, Yves Van de Peer, Alasdair J Nisbet

## Abstract

Psoroptic mange, caused by infestation with the ectoparasitic mite, *Psoroptes ovis*, is highly contagious, resulting in intense pruritus and represents a major welfare and economic concern for the livestock industry Worldwide. Control relies on injectable endectocides and organophosphate dips, but concerns over residues, environmental contamination, and the development of resistance threaten the sustainability of this approach, highlighting interest in alternative control methods. However, development of vaccines and identification of chemotherapeutic targets is hampered by the lack of *P. ovis* transcriptomic and genomic resources. Building on the recent publication of the *P. ovis* draft genome, here we present a genomic analysis and transcriptomic atlas of gene expression in *P. ovis* revealing feeding- and stage-specific patterns of gene expression, including novel multigene families and allergens. Network-based clustering revealed 14 gene clusters demonstrating either single- or multi-stage specific gene expression patterns, with 3,075 female-specific, 890 male-specific and 112, 217 and 526 transcripts showing larval, protonymph and tritonymph specific-expression, respectively. Detailed analysis of *P. ovis* allergens revealed stage-specific patterns of allergen gene expression, many of which were also enriched in “fed” mites and tritonymphs, highlighting an important feeding-related allergenicity in this developmental stage. Pair-wise analysis of differential expression between life-cycle stages identified patterns of sex-biased gene expression and also identified novel *P. ovis* multigene families including known allergens and novel genes with high levels of stage-specific expression. The genomic and transcriptomic atlas described here represents a unique resource for the acarid-research community, whilst the OrcAE platform makes this freely available, facilitating further community-led curation of the draft *P. ovis* genome.

## Background

Psoroptic mange, caused by the ectoparasitic mite *Psoroptes ovis*, is characterised by pruritus and skin irritation and is a major welfare and economic concern for the livestock industry as the parasite infests both cattle and sheep, causing the disease “sheep scab” in the latter [1,2]. In sheep, control relies on injectable macrocyclic lactone-based endectocides and organophosphate dips but concerns over residues, environmental contamination and the development of resistance threaten the sustainability of this approach and have highlighted interest in developing alternative control methods [3,4]. However, the development of novel interventions (including vaccines and the identification of potential chemotherapeutic targets) has previously been hampered by a lack of detailed transcriptomic and genomic resources for *P. ovis*.

The integration of newly-available transcriptomic and genomic data with current knowledge of the basic biology of the mite is pivotal in the development of such novel interventions: The basic biology of the obligate ectoparasitic mite, *P. ovis*, on sheep is well understood, with the life-cycle taking place entirely on the ovine host and lasting from 11-19 days from egg hatch to egg production by the adult [5]. The life-cycle progresses from egg through four developmental stages (larvae → protonymph → tritonymph → adult (male/female)) (Figure 1). Adult female mites can survive on the host for up to 42 days and during this time they may deposit up to 80 eggs [1,6,7]. *Psoroptes ovis* mites are able to survive for a limited time (15-16 days) off-host, enabling their transfer from animal to animal via fomites [8]. *Psoroptes ovis* is a non-burrowing mite, which feeds at the skin surface consuming serous exudate, lymph and red blood cells [9]. Mites survive on the surface of the skin and their mouthparts, which are thought to abrade rather than pierce the skin, do not penetrate beyond the stratum corneum, the outermost layer of the skin [10]. As the mites move across the surface of the skin they secrete and excrete allergens and other potent pro-inflammatory factors and this combination of mechanical skin abrasion, allergen deposition and grooming behaviour by the host in response to the pruritus caused by the mites all contribute to the subsequent cutaneous inflammatory response [11–13]. However, the role of the different developmental stages of *P. ovis* in eliciting the pathology associated with the host pro-inflammatory response, and subsequent semi-protective immunity, is currently unknown and would be greatly improved with knowledge of the individual life-cycle stage transcriptomes.

**Figure 1.**
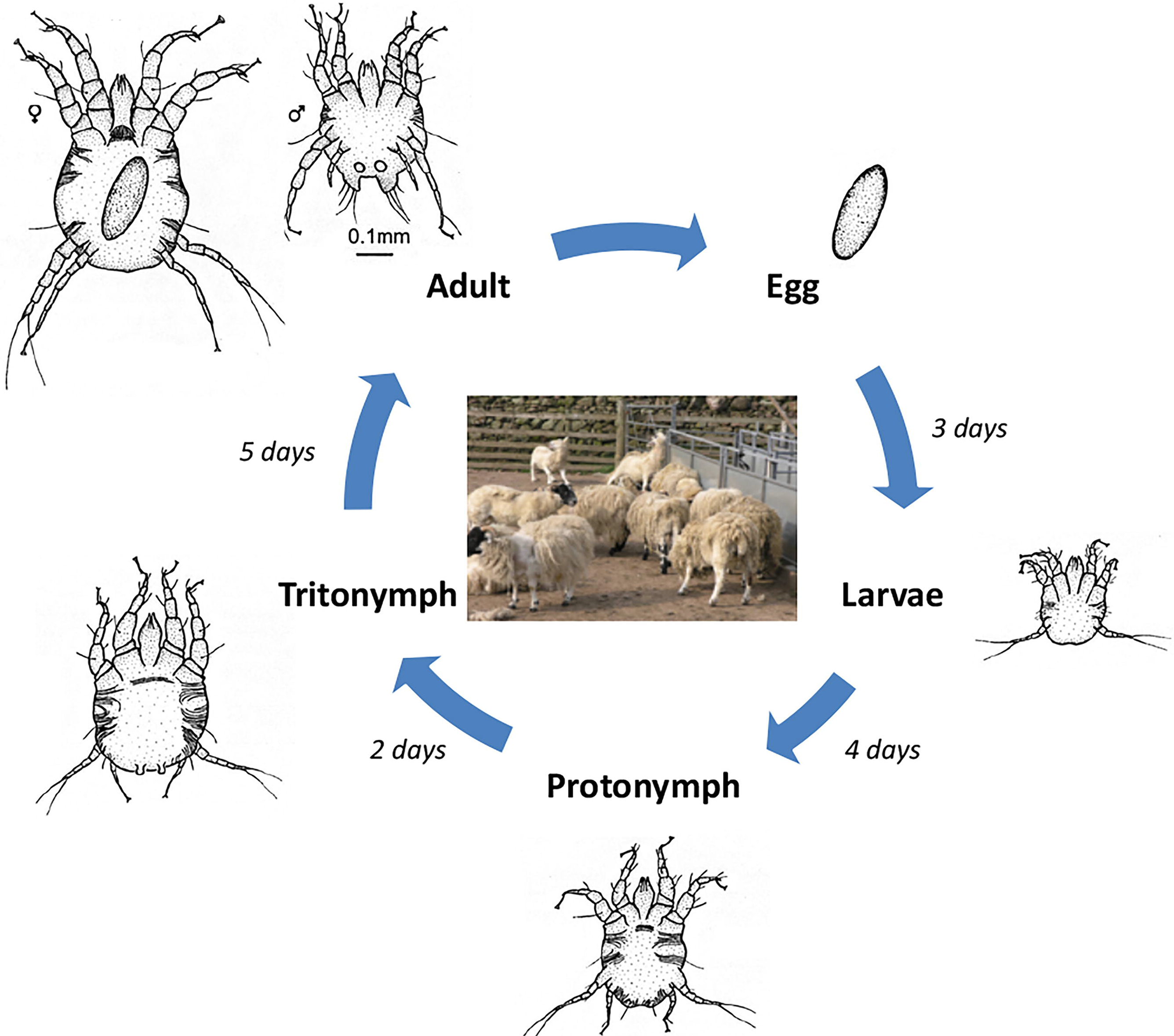
*Psoroptes ovis* life-cycle. Image demonstrates progression from egg, through larvae (L), nymph stages (protonymph (P) and tritonymph (T)) and onto adult male (AM) and adult female (AF). Image adapted from “Diagram of the life-cycle of *Psoroptes ovis* parasitic mite of sheep and cattle” (commons.wikimedia.org/wiki/File:Life-cycle-psoroptes-ovis-mite-diagram.jpg) under Creative Commons License (CC-BY-SA-3.0).

Existing transcriptomic tools and resources for *P. ovis* are limited and include an expressed sequence tag (EST) survey of ~500 *P. ovis* cDNAs [14], a subtractive suppressive hybridisation (SSH) based comparison of gene expression between “fed” and “starved” *P. ovis* mites [15] and a cDNA microarray based on ~1000 *P. ovis* ESTs [16]. More recently a preliminary transcriptomic analysis of *P. ovis* var. *cuniculi* across a limited number of developmental stage comparisons using Illumina RNA-seq was described [17]. The recent generation of the *P. ovis* genome, which included the prediction and annotation of the *P. ovis* transcriptome [18] has substantially improved the resources available and enables more detailed genomic and transcriptomic analyses of *P. ovis*. At 63.2Mb, the draft genome assembly demonstrated that *P. ovis* has one of the smallest arthropod genomes sequenced to date, smaller than the genome of the two-spotted spider mite (*Tetranychus urticae* (90Mb)) but comparable in size with the closely related house dust mite (HDM) genomes (*Dermatophagoides farinae* (53.5Mb) and *D. pteronyssinus* (70.76Mb)) and the ectoparasitic scabies mite (*Sarcoptes scabiei* (56.2Mb)) [18–22]. Herein, using the recently described *P. ovis* genome [18], we described the detailed annotation of the genome to Gene Ontology (GO) level along with a quantitative transcriptomic analysis of *P. ovis* gene expression across multiple life-cycle stages, providing for the first time a complete transcriptomic atlas of stage-specific and feeding-related gene expression in this economically-important ectoparasite of livestock.

## Results and Discussion

### Functional annotation of the P. ovis predicted transcriptome derived from the draft genome

Overall, 12,041 predicted protein coding genes were identified in the *P. ovis* genome, which represented the first global survey of the *P. ovis* gene repertoire [18]. This represents ~190 genes per Mb for *P. ovis*, which is comparable to other closely related mite species, for example *T. urticae* (205 genes per Mb), *S. scabiei* (189 genes per Mb), *D. farinae* (306 genes per Mb) and *D. pteronyssinus* (177 genes per Mb). Interproscan analysis resulted in further functional annotation for 9,960 genes and significant BLAST hits against the National Center for Biotechnology Information (NCBI) non-redundant (nr) database (March 2018) were identified for 10,009 (83%) genes. Gene ontology (GO) assessment was performed in Blast2GO resulting in the assignment of GO terms for 8,681 (72%) genes and functional annotation for 7,614 (63%) genes. Figure 2 shows the distribution of sequences per GO term across multiple classification levels and is presented as three pie-charts showing GO term distributions for Biological Process, Molecular Function and Cellular Component. GO terms for Biological Process were further subdivided between 13 categories, including: cellular macromolecule biosynthetic process (12%), RNA metabolic process (11%), phosphate-containing compound metabolic process (11%) and signal transduction (10%). The Molecular Function category was further split into 7 subcategories including: hydrolase activity (21%), transferase activity (18%), protein binding (17%), nucleic acid binding (15%) and catalytic activity, acting on a protein (13%). GO terms for Cellular Component were subdivided into 6 categories, including: intracellular organelle part (20%); cytoplasmic part (19%) and protein-containing complex (19%).

**Figure 2.**
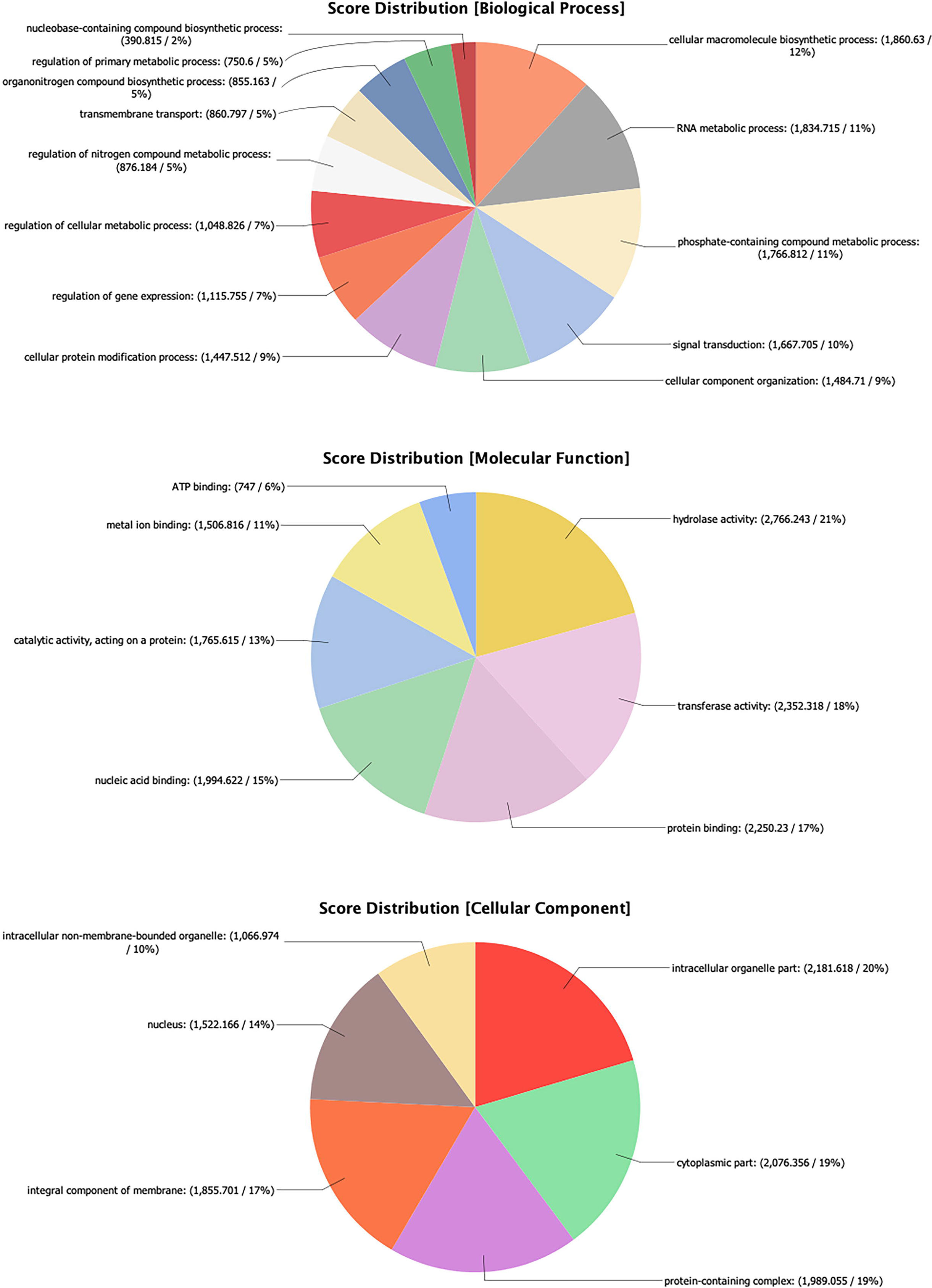
*Psoroptes ovis* genome Gene Ontology (GO) annotation Each chart shows the multilevel distribution of sequences per GO term. Distribution of GO terms are summarised across three main categories: Biological Process, Molecular Function and Cellular Component.

### Interactive web-based presentation of the entire *P. ovis* genome and gene expression atlas allows interrogation of individual genes and their stage-specific expression profiles

The full annotation of the *P. ovis* genome has now been made publicly available via the Online Resource for Community Annotation of Eukaryotes (OrcAE) and to maximise the utility of this information for researchers, for each gene we created a gene-specific page, describing the full annotation available for that gene, including information relating to: gene function, GO terms, Pfam protein domains, protein homologues and significant BLAST hit data, gene structure, coding sequence, protein sequence and, where available, transcript evidence based on associated ESTs/cDNA data (Figure 3). This gene expression atlas also features a fully searchable database of the entire genome assembly as well as incorporating a visual and numerical display of the gene expression data (RNA-seq) across the *P. ovis* life-cycle stages, allowing full interrogation of expression profiles for target genes of interest. Each gene was assigned a unique loci identifier with the following format: psoviXXgYYYYY, where XX defines the scaffold ID and YYYYY denotes the specific location within the scaffold. As with most large-scale genome projects, the *P. ovis* genome relied upon a computational gene prediction and annotation pipeline and although we also incorporated additional RNA-seq data as transcript evidence in this process it is likely that some errors will remain. As such continual manual curation of gene prediction and annotation remains a critical step in assessing and improving gene prediction accuracy and overall confidence in the genome. It is clear that incorrect and incomplete annotations have the capacity to tarnish each subsequent experiment that relies on them, making the provision of accurate and up-to-date annotation essential [23]. Therefore, continual manual curation of genes and associated gene families by experts in the field will further enhance the value of gene predictions for the entire research community. To facilitate this, the OrcAE platform is unique in that it also provides the tools and information for community-led manual validation of gene annotations [24]. The system is built on a Wiki philosophy, meaning that all modifications to a certain gene are stored and logged in the gene history. In order to be able to modify genes, users must apply for an editable account and we encourage the acarid research community to participate in this process. However, anonymous users are free to browse the publicly available *P. ovis* genome but will not have editing rights.

**Figure 3.**
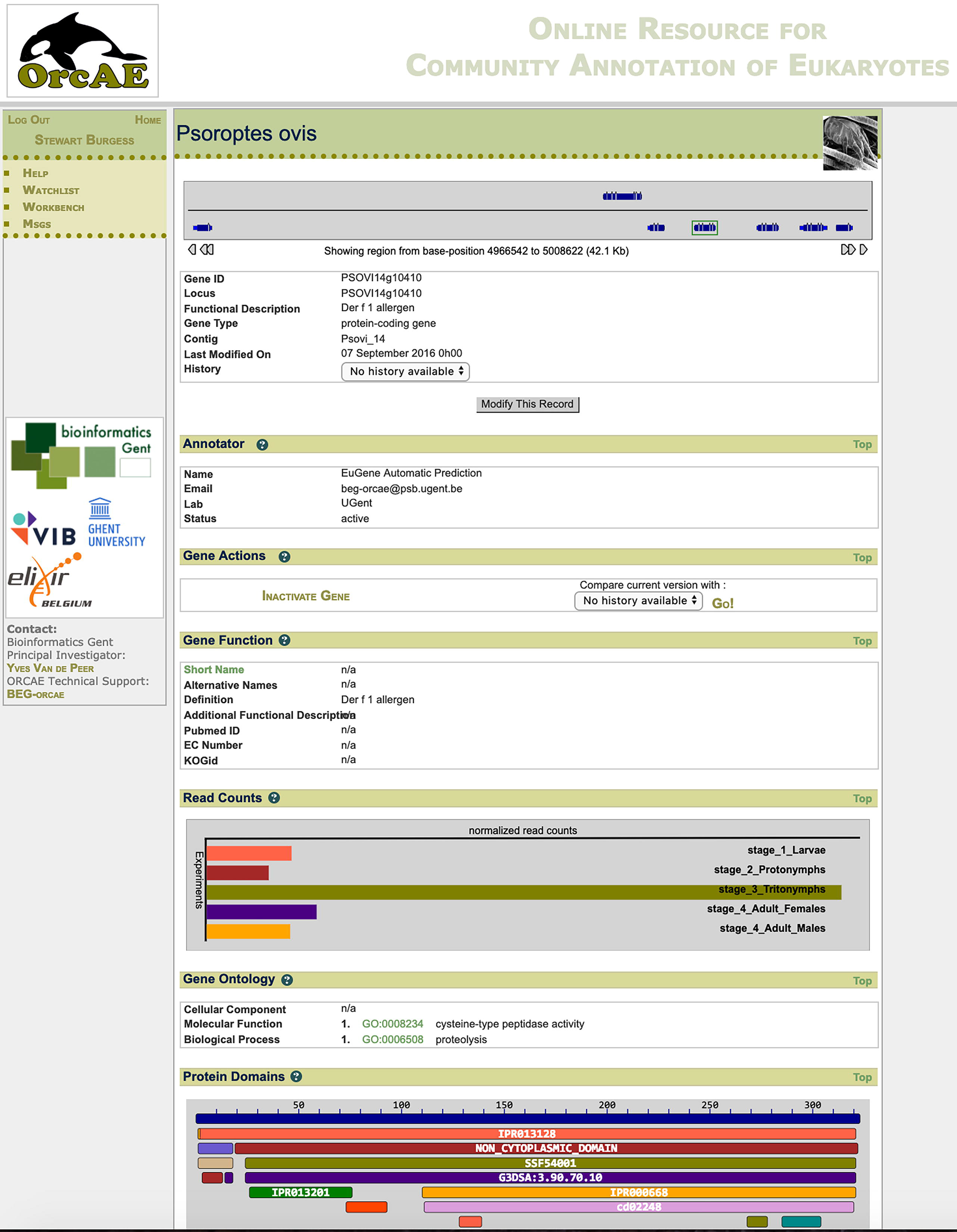
Example page of the web-based *P. ovis* gene expression atlas. The atlas was constructed within the Online Resource for Community Annotation of Eukaryotes (OrcAE) framework [24]. Here we show the gene-specific page for the major *P. ovis* allergen, Pso o 1 (psovi14g10410). Website: http://bioinformatics.psb.ugent.be/orcae/overview/Psovi

### P. ovis collection, life-cycle staging and preparation of “fed” and “starved” mite populations

Staging of *P. ovis* into individual life-cycle stages (Figure 1) is challenging and time-consuming but can be performed with a high degree of accuracy [25]. Whilst staging of adult males (length: 396μm, width: 380μm) and females (length: 536μm, width: 467μm) was relatively straightforward by size alone, staging of larvae relied on identification by size (length: 250μm, width: 212μm) and the presence of three pairs of legs (rather than the four found in nymphs and adults). The relatively small size of larvae also required the collection of high numbers of individuals to allow recovery of sufficient RNA for sequencing. Separation of the nymph stages: protonymphs (length: 309-313μm, width: 292-351μm) and tritonymphs (length: 402-414μm, width: 370-436μm) was partially achieved by size, but also relied on the identification of key morphological differences as highlighted in the taxonomical key provided by Sanders *et al* [25]. In total 1,479 Adult Females, 1,618 Adult Males, 3,676 Larvae, 1,814 Protonymphs and 1,194 Tritonymphs were individually staged and collected. The “fed” and “starved” *P. ovis* mite samples (n=3/each) were taken from the same mixed population, prior to staging and split into 3 pools.

### RNA extraction and Quality Control

For each life-cycle stage and for the “fed” and “starved” conditions, the mites were divided into three equal-sized pools and high quality RNA was extracted from each pool, yielding >10μg total RNA (5μg of which was used for the generation of each RNA-seq library) and RNA integrity numbers (RIN values) of greater >7.5 were obtained for each sample.

### RNA-seq profiling of *P. ovis* stage-specific and feeding-related gene expression

Illumina sequencing resulted in 8-26 million raw sequence reads for each of the twenty one sequencing libraries (three biological replicates for each of the five life-cycle stages and three each for “fed” and “starved” mites) with a mean of 13.7 million reads per sample (Table S1). For each replicate, from each developmental stage we generated a set of expression estimates from the trimmed reads, as transcripts per million (TPM) using the transcript quantification tool Kallisto (Version 0.44.0 [26]) and the predicted transcriptome derived from the *P. ovis* genome [18].

### Network analysis and clustering of stage-enriched gene expression in P. ovis

In order to identify groups of genes whose expression is associated with either single- or multiple-developmental stages of *P. ovis*, we performed a network graph analysis. It is well characterised that genes playing distinct roles in common signalling pathways or biological processes often share similar patterns of expression and therefore regulation [27]. As such, when genes are found to have similar expression profiles across multiple samples or sample classes, i.e. they are co-expressed, this may be an indication that they share functional or biological activity, i.e. guilt-by-association/guilt-by-profiling [28]. Read count data, expressed as TPM, for each replicate, from each life-cycle stage was used to generate a gene-gene network graph within the Graphia Professional package [29]. The network graph was generated using a Pearson correlation cut-off of 0.9, resulting in a graph with 10,655 nodes (genes) connected by 3,451,719 edges. Individual genes were clustered within the Graphia Professional package using a Markov Cluster Algorithm (MCL) inflation value of 2.2, resulting in a final gene-to-gene network graph consisting of 7,719 genes divided across 50 clusters (Figure 4). The resulting network graph consisted of nodes (genes) connected by virtue of the similarity of their gene expression profile across each of the *P. ovis* life-cycle stages (Figure 4).

**Figure 4.**
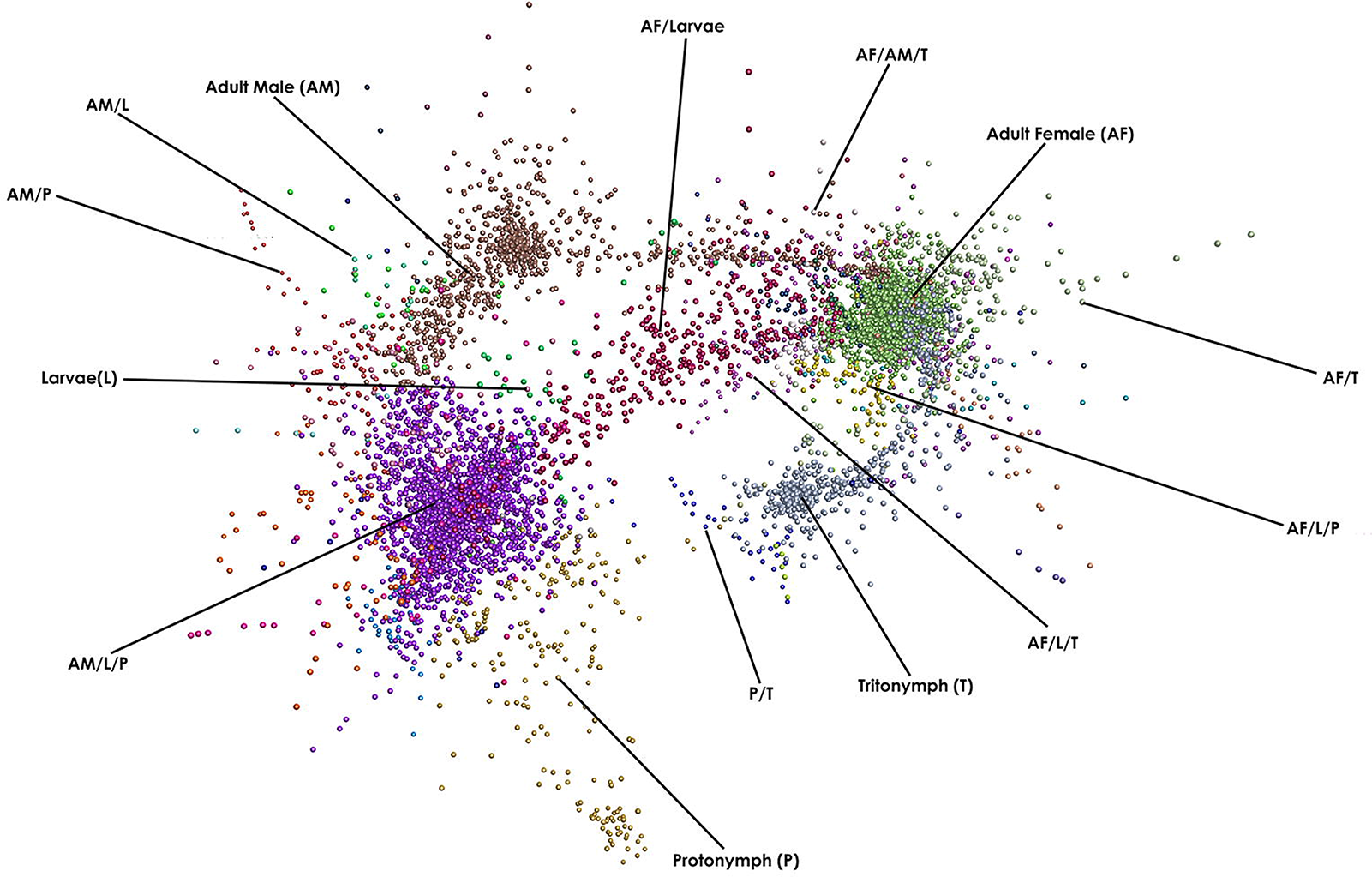
Network clustering of *P. ovis* stage-specific gene expression profiles. A Pearson correlation matrix was generated comparing the gene expression data derived from all of the *P. ovis* life-cycle stages. A network graph was constructed using a Pearson correlation cut off value of 0.9. Each node (coloured circle) represents a single *P. ovis* gene. The network graph was clustered using an MCL inflation value of 2.2 with each cluster of genes being represented by a different colour. Clusters of genes exhibiting expression profiles specific to either single- or multiple-developmental stages of *P. ovis* are labelled as follows: Larvae (L), Protonymph (P), Tritonymph (T), Adult Female (AF) and Adult Male (AM).

Amongst the 50 gene clusters within the network, a number of clusters demonstrated similar patterns of expression across the *P. ovis* life-cycle stages and these were further collated into groups of clusters. This resulted in a final total of 14 gene clusters (Table 1) demonstrating either single- or multi-stage enriched gene expression patterns (Figure 5). The genes attributed to each individual cluster are listed in Supplementary File 1.

**Table 1.**
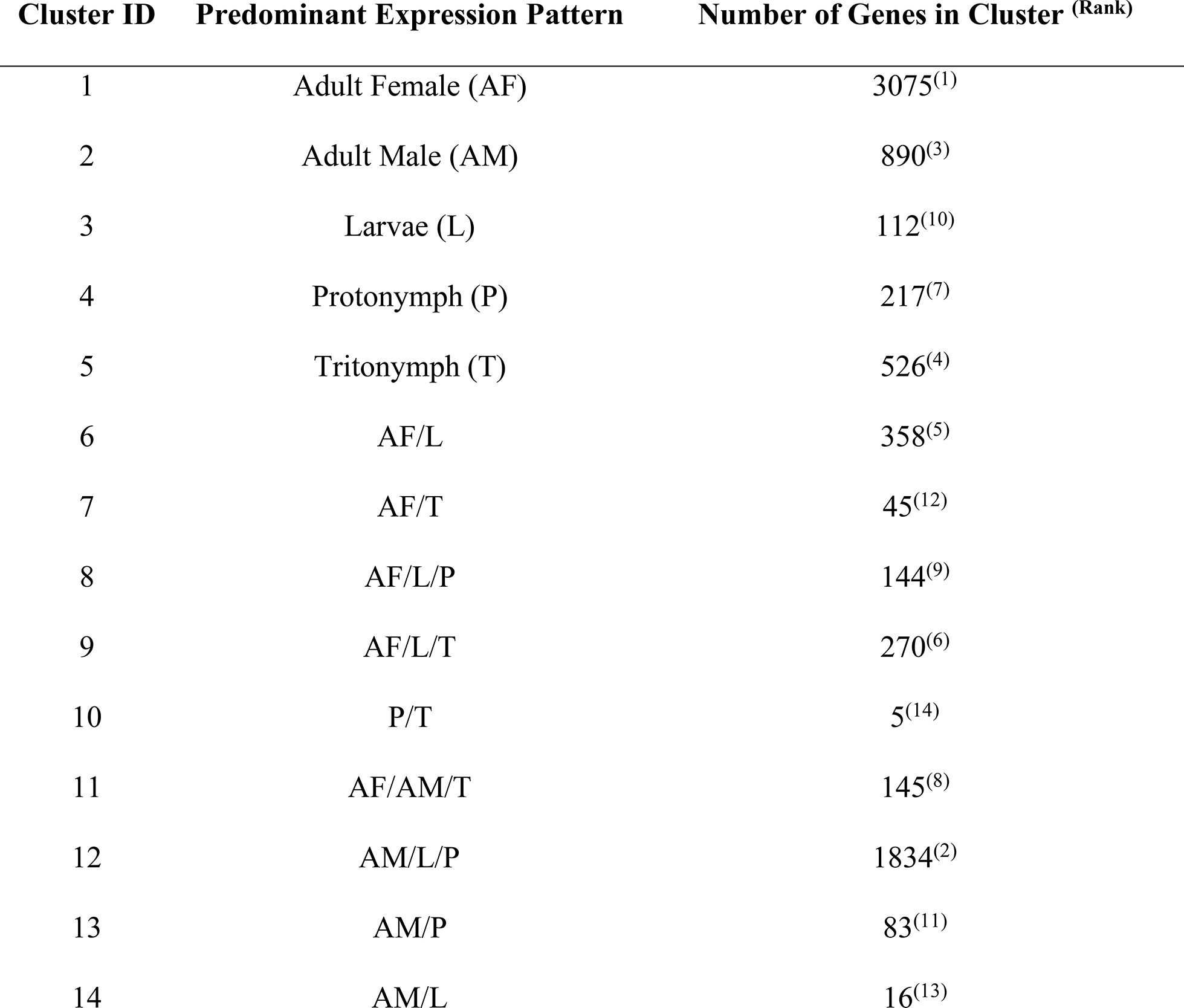
Description of final *P. ovis* stage-specific gene expression clusters (n=14).

**Figure 5.**
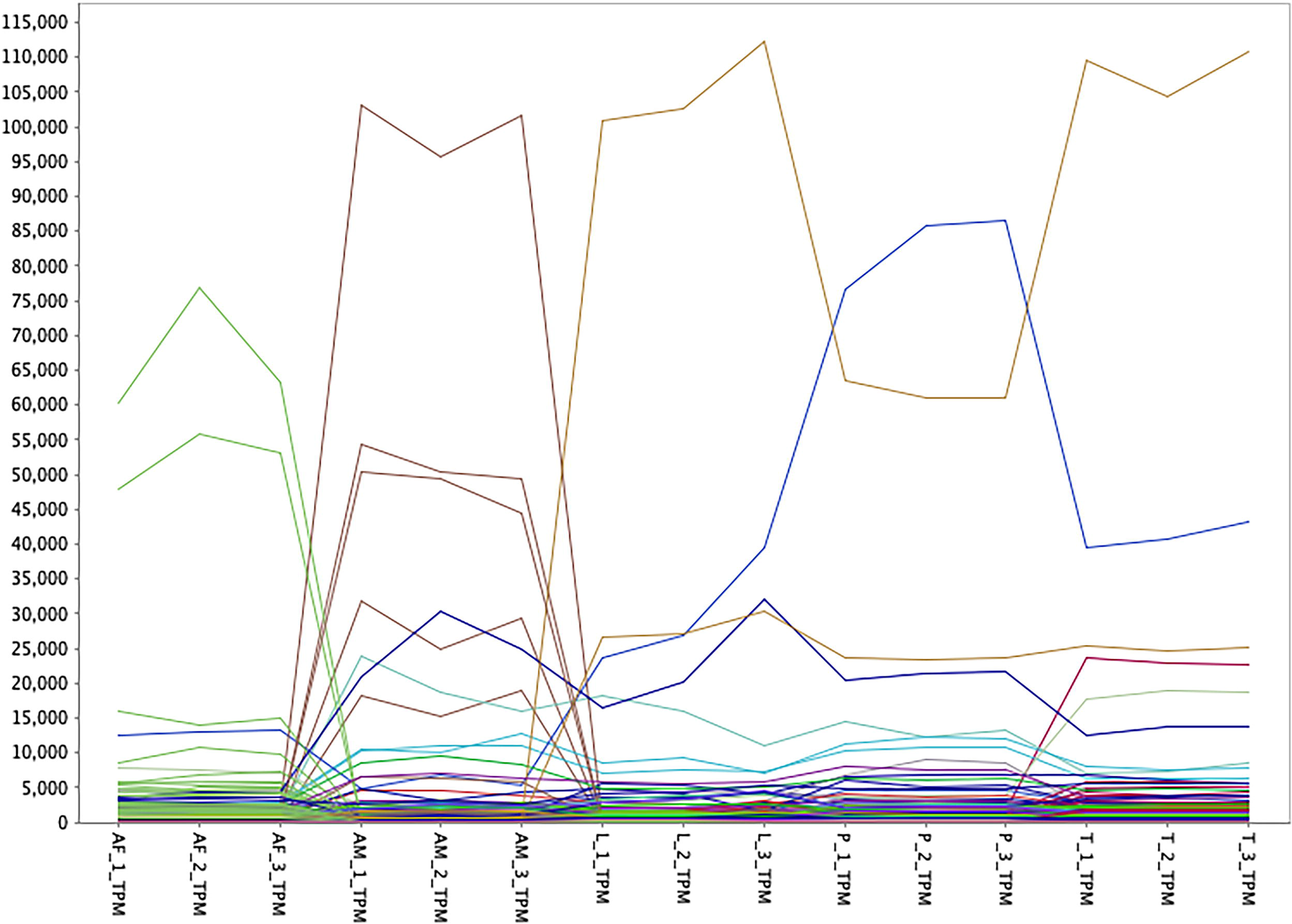
Line plot showing *P. ovis* stage-specific gene expression profiles. Each line represents the expression profile of a single gene across triplicate samples for each *P. ovis* life-cycle stage: colours indicate individual gene clusters: Adult Female (AF), Adult Male (AM), Larvae (L), Protonymph (P), Tritonymph (T). Y-axis shows read count values expressed as transcripts per million (TPM) as derived from Kallisto (Version 0.44.0 [26]).

### Functional annotation of *P. ovis* stage-specific gene expression clusters

The genes within each cluster were then mapped back to the original *P. ovis* genome annotation [18] and a Gene Ontology (GO) analysis was performed within the Blast2GO package to identify associated GO terms for molecular function, biological process and cellular component attributed to each cluster.

### Genes enriched in *P. ovis* adult females (AF) - Cluster 1

Similar to other mite species, *P. ovis* exhibits sexual dimorphism, with clear morphological and behavioral differences existing between adult males and females and potentially also during the tritonymph stage [5]. In the fruit fly *Drosophila melanogaster*, many of these sex-related changes have been attributed to differences in gene expression, suggesting that sexual dimorphism may result primarily from the differential expression of genes present in both sexes, e.g. sex-biased gene expression [30–32]. Here we investigated the differential expression of genes between adult male and adult female *P. ovis* mites in order to elucidate the mechanisms underlying this sexual dimorphism. Clustering analysis identified a cluster of 3,075 genes showing adult female-enriched patterns of gene expression, which represents the largest cluster of stage-specific genes identified in this study. Amongst these, genes encoding *P. ovis* homologues of vitellogenin (psovi09g01710), vitellogenin receptor (psovi63g00310) and two group 14 apolipophorin allergens (psovi73g00070 and psovi35g00110) were present. Each of these genes has a role in oogenesis and all showed almost exclusive expression in the female mites. In addition, a number of genes encoding proteins involved in lipid metabolism within the same cluster, including an alkylglycerol monooxygenase-like protein (psovi288g00950), a perilipin-like protein (psovi66g00530), an ATP-binding cassette transporter (ABCA1) lipid exporter protein (psovi284g05730) were present along with 14 genes encoding proteins involved in the elongation of long chain fatty acids. The expression of genes involved in lipid processing pathways in the adult females may indicate a role in nutrition related to supply of nutrients to the reproductive tissues. Lipids represent a major constituent of the sheep epidermis and *P. ovis* is likely to use the abundant supply of lipids as an energy-rich food source [13,33,34]. One of the most highly expressed female-enriched genes encodes a putative serine protease inhibitor (Serpin) leukocyte elastase inhibitor-like protein (psovi22g04610) homologous to both the newly-characterised HDM Der f 27 allergen and to a scabies mite (*S. scabiei*) serpin SMSB4, which has been shown to interfere with host complement-mediated neutrophil functions and promote staphylococcal growth during infestation with *S. scabiei* [35,36]. A further four uncharacterised and highly expressed genes (psovi09g01110, psovi72g00350, psovi08g00890 and psovi06g00980) showed almost exclusive expression in the female mites. However, no significant BLAST hits were identified for these genes, indicating that they may represent unique *P. ovis* genes of as yet unknown function. A further female-enriched gene with a high level of expression in females (psovi14g01150) showed a significant BLAST hit against a skin secretory protein, xP2-like from the African clawed frog, *Xenopus laevis* and the scallop *Mizuhopecten yessoensis*. This gene encodes a protein with a Trefoil (P-type) domain, which is a cysteine-rich domain consisting of approximately 45 amino-acids found in selected extracellular eukaryotic proteins [37]. The Trefoil domain has been identified as one of a relatively small number of protein families that represent potential allergens [38], as such the nature of this *P. ovis* gene and its role in the pro-inflammatory/allergic type response in psoroptic mange warrants further investigation. A final group of genes co-located in the female-enriched cluster are putative heat shock proteins (Hsp) with ten genes representing potential Hsps including 10kDa, 60kDa, 70kDa, 83kDa and 90kDa protein encoding genes. Hsp70 has been identified as an important allergen from sesame seed and hazel pollen [39] and has also been shown to be upregulated in the honey bee parasitic mite, *Varroa destructor* following thermal stress and exposure to acaricide, indicating a potential role in xenobiotic metabolism [40].

### Genes enriched in *P. ovis* adult males (AM) - Cluster 2

Network clustering analysis identified 890 genes showing an adult male-enriched pattern of gene expression. Many of the genes in this cluster play a role in muscle development and function including genes encoding paxillins, muscle LIM proteins, PDZ and LIM domain containing proteins, alpha actinin, microtubule actin cross-linking factor, vinexins, paramyosin, vinculin, titin and calmodulin. Many of these genes are involved in the formation and contraction of muscle fibres and this may be a reflection of the increased motility of *P. ovis* males, compared to adult female mites [41]. This cluster also contained 19 highly expressed genes with an average read count of >1000 in the males, many of which currently represent uncharacterised genes with no known homologues. Perhaps the most interesting of these are a group of three genes, co-located on the largest scaffold (Psovi_22) of the *P. ovis* genome (psovi22g004350; psovi22g004360 & psovi22g004380) all of which are highly expressed in male mites. All three genes encode short proteins of ~19kDa (166-169 amino acids) two of which (psovi22g004360 & psovi22g004380) share 69% identity at the amino acid level. In part, the presence of these highly expressed genes within potential multigene families may be explained by previous studies in *D. melanogaster*, where it has been shown that genes demonstrating a male-biased pattern of expression often exhibit greater differences in gene expression levels than that observed for female-biased genes or for non-sex-biased genes [42,43]. In addition, the rates of evolution at the sequence level observed in sex and reproduction-related (SRR) and non-SRR genes in *D. melanogaster* also differs, with male and female SRR genes evolving more rapidly than non-SRR genes [44–47]. In addition, Connallon and Knowles [30] showed that the majority of this sex-biased gene expression was due to adaptive changes in the males, suggesting they may experience stronger selection pressures than the females.

### Genes enriched in *P. ovis* larvae (L) - Cluster 3

The cluster of larvae-enriched genes represents one of the smallest single-stage clusters with only 112 genes. The most highly expressed gene in the cluster showed very similar levels of expression in both larvae and tritonymphs and shares significant homology with a novel gene of unknown function from the HDM *D. farinae* [48]. The protein encoded by this gene was termed *Dermatophagoides farinae* most abundant protein 2 (DFP2) and as yet remains uncharacterised functionally [48]. Two further highly expressed genes in this cluster encode *P. ovis* homologues of an ADP/ATP translocase (psovi43g01550) and a chaperonin containing TCP1 complex protein (CCT1) (psovi283g00230). CCT1 forms part of a chaperonin complex consisting of two identical stacked rings, each containing eight different proteins [49]. The completed complex functions in an ATP-dependent manner and is responsible for folding a range of proteins, including actin and tubulin [50].

### Genes enriched in *P. ovis* protonymphs (P) - Cluster 4

Clustering analysis identified a cluster of 217 genes showing a protonymph-enriched pattern of gene expression. Seventeen genes in this cluster shared significant homology with the HDM, DFP2 gene, as described above. However, the present study shows, for the first time, that multiple copies of this gene may exist in the *P. ovis* genome and that many of these are highly expressed across the juvenile stages (including, larvae, protonymphs and tritonymphs). The GO analysis performed within the Blast2GO package demonstrated that, for the protonymph-enriched cluster, 22% of genes in the Biological Process category were associated with chitin metabolic processes, whilst 13% of genes in the Molecular Function category are involved in chitin binding, equating to 22 and 14 genes, respectively. Of these, specific functional annotation for chitin binding and metabolism was available for 12 genes, including a putative *P. ovis* chitin synthase (psovi294g01310), a chitin deacetylase (psovi33g00850), five chitinase homologues (psovi14g06600, psovi05g01800, psovi14g09170, psovi283g03530 and psovi20g01930) and five chitin binding peritrophin-like genes (psovi14g06590, psovi14g10800, psovi14g06580, psovi286g01240 and psovi284g00080). The same cluster also contained seven genes encoding putative cuticle proteins, all of which are highly expressed across the juvenile stages and not just the protonymphs. The presence of many highly expressed cuticle and chitin binding transcripts across the juvenile stages, indicates a role in the formation and/or moulting of the mite cuticle. This cluster also contained a further putative serpin with homology to the SMSB3 serine protease inhibitor from *S. scabiei* (psovi22g04600) which has a role in inhibition of host complement during scabies mite infestation [51]. Interestingly, this gene is co-located adjacent to the female-specific serpin described above (psovi22g04610) with the predicted proteins from these two genes sharing ~30% identity at the amino acid level. A further cluster (Cluster 10) contained five genes with an expression profile enriched in both protonymphs and tritonymphs and these represent two further DFP2 homologues, two putative cuticle proteins and one uncharacterised gene.

### Genes enriched in *P. ovis* tritonymphs (T) - Cluster 5

Clustering analysis identified a cluster of 526 genes demonstrating tritonymph-enriched patterns of gene expression. GO analysis showed that for the tritonymph-enriched cluster, a large proportion (53%) of the genes in the Molecular Function category were involved in enzyme activities, including proteolysis (8%), transferase activity (12%), hydrolase activity (20%) and oxidoreductase activity (13%). A similar pattern was observed for the Biological Process category with 16% of genes involved in oxidation-reduction processes and 10% in proteolysis. In total, 73 of the 526 genes in this cluster (14%) were assigned Enzyme Commission (EC) codes, distributed as follows: hydrolases (n=27), oxidoreductases (n=21), transferases (n=15), lyases (n=5), ligases (n=4) and isomerases (n=1). A further factor contributing to the large numbers of enzymes within this cluster could be the presence of numerous putative *P. ovis* allergens, many of which are homologues of HDM allergens and several of which show protease activity [52]. Putative allergen genes enriched within the tritonymph cluster included some of the most abundant *P. ovis* allergens, i.e. Pso o 1, 2, 7, 8, 13 and 21, along with Pso o 18, 30, 34 and 36. As we observed in the larvae and protonymph enriched clusters, a number of DFP2 homologues (n=17) were also enriched within the tritonymph cluster, although the expression levels for these transcripts were often lower than that observed in the protonymph stage. A further group of genes (n=12) enriched in the tritonymph cluster belong to an as yet functionally-uncharacterised group of senescence-associated proteins from the two-spotted spider mite, *T. urticae* [20]. Another group of five tritonymph-enriched genes is co-located on a single scaffold of the *P. ovis* genome (psovi22g03270, psovi22g03310, psovi22g03330, psovi22g03320, psovi22g03340) indicating a potential shared expression and function profile. These genes showed high levels of expression in the tritonymphs but no known homologues were found from the BLAST searches, indicating that these may be unique *P. ovis* genes.

### Genes enriched in *P. ovis* adult females (AF), larvae (L) and tritonymphs (T) - Cluster 9

The cluster of genes with similar patterns of expression across *P. ovis* adult females, larvae and tritonymphs contained 270 genes. GO analysis showed that 47% of those within the Molecular Function category were involved in either RNA-binding (21%) or as structural constituents of ribosomes (26%). A similar picture was observed in the Biological Process category with 32% involved in translation and 18% in ribosome biogenesis. A closer look at the transcripts involved revealed that 54 of the 270 genes (20%) showed homology with 40S or 60S ribosomal protein genes many of which were amongst the most highly expressed genes in the cluster.

### Genes enriched in *P. ovis* adult males (AM) and protonymphs (P) - Cluster 13

Similar to the adult male-enriched cluster, many of the most highly expressed genes in the adult male/protonymph-enriched cluster encode proteins involved in formation and contraction of muscle tissues, including two homologues of a *D. melanogaster* muscle-specific protein 20-like (psovi284g04960 and psovi43g00510), the *P. ovis* allergens tropomyosin (Pso o 10) and paramyosin (Pso o 11), three troponin homologues (troponin-I, -C and -T), five myosin genes (heavy and light chain) and a *P. ovis* arginine kinase (psovi292g02430). As with the male-enriched cluster the large number of muscle-specific transcripts may be a reflection of the increased motility of the *P. ovis* males and protonymph stages [41].

### Assessment of the most abundantly expressed genes for each life-cycle stage

For each of the *P. ovis* life-cycle stages we identified the top 100 most abundantly expressed genes (i.e. highest TPM value). To assess the expression of these genes across the different life-cycle stages we used a five-way Venn/Euler diagram to examine their expression across *P. ovis* life-cycle stages (Figure 6). The IDs and annotations of the genes attributed to each arm of the Venn diagram are detailed in Supplementary File 2. As can be seen in Figure 6, a number of the most abundantly expressed genes demonstrated stage-enriched expression, for example 43 genes were enriched in females, 21 in males, 14 in tritonymphs, 11 in larvae and 6 in protonymphs. The female-enriched gene cluster included both vitellogenin and the vitellogenin receptor and a homologue of the recently classified HDM allergen, Der f 27 (a serine protease inhibitor (serpin)) termed here Pso o 27, which showed very high levels of expression in female mites. We also identified an ABCA1 homologue, which may have a role in lipid transport and more specifically in the removal of cholesterol from cells [53–55]. The cluster of 14 genes abundantly expressed in tritonymphs included a number of allergens, notably two copies of the major *P. ovis* allergen Pso o 1 (psovi14g10410 & psovi14g10420) a homologue of the HDM allergen Der p 1, which showed expression across all life-cycle stages but with significantly (p≤0.05) higher expression in tritonymphs [56]. In addition, the cluster contained both copies (psovi17g08010 & psovi88g00180) of a Group 2 (Der p 2) allergen homologue (Pso o 2), which is a potential functional mimic of the TLR4 accessory protein MD-2 and may play a role in host immune activation [57] and is also used as a diagnostic antigen for the detection of sheep scab [58]. The same cluster also contained a Group 21 (Der p 21) allergen homologue, termed Pso o 21, which has been shown to trigger IL-8 production in airway epithelial cells through a TLR2-dependent mechanism [59] and a Group 14 (Der f 14 or M-177) apolipophorin-like allergen homologue, termed Pso o 14, characterised as a large lipid binding protein with IgE-binding and cytokine-inducing capacities [60]. Insect apolipophorins have also been demonstrated to have a role in pattern recognition as part of the insect innate immune response to fungal beta-1,3-glucan [61]. In addition, 37 of the most abundantly expressed genes were conserved across stages (Figure 6). Of these conserved abundant genes many are involved in the formation and contraction of muscle tissues (i.e. actin, tropomyosin, troponin, arginine kinase, muscle LIM protein 1 and smoothelin) or in glycolysis and energy metabolism (glyceraldehyde-3-phosphate dehydrogenase, fructose-bisphosphate aldolase, cytochrome c oxidase subunits II and III, ATP synthase F0 subunit 6 and ADP/ATP translocase) supporting the central importance of these key biological processes across all *P. ovis* life-cycle stages. One of these gene products, fructose-bisphosphate aldolase, has been identified as a potential allergen in the red shrimp (*Solenocera melantho*) [62] and this may represent a novel mite allergen. Interestingly, genes with similar central functions were also found in a cluster of 19 genes conserved between larvae, protonymphs, tritonymphs and males but were not in the top 100 most abundantly expressed genes in females, for example NADH dehydrogenase subunit 6, arginine kinase, troponin, muscle LIM protein 1 and smoothelin. The same cluster also contained a putative cuticle protein, a putative secreted salivary gland protein and a number of myosin light chain genes. In addition, 10 genes were conserved between juvenile stages (larvae, protonymphs and tritonymphs) including 3 genes encoding homologues of a *D. farinae* DFP2-like protein of unknown function, a putative cuticle protein and a mu-class glutathione-S-transferase (GST) or Group 8 HDM allergen homologue.

**Figure 6.**
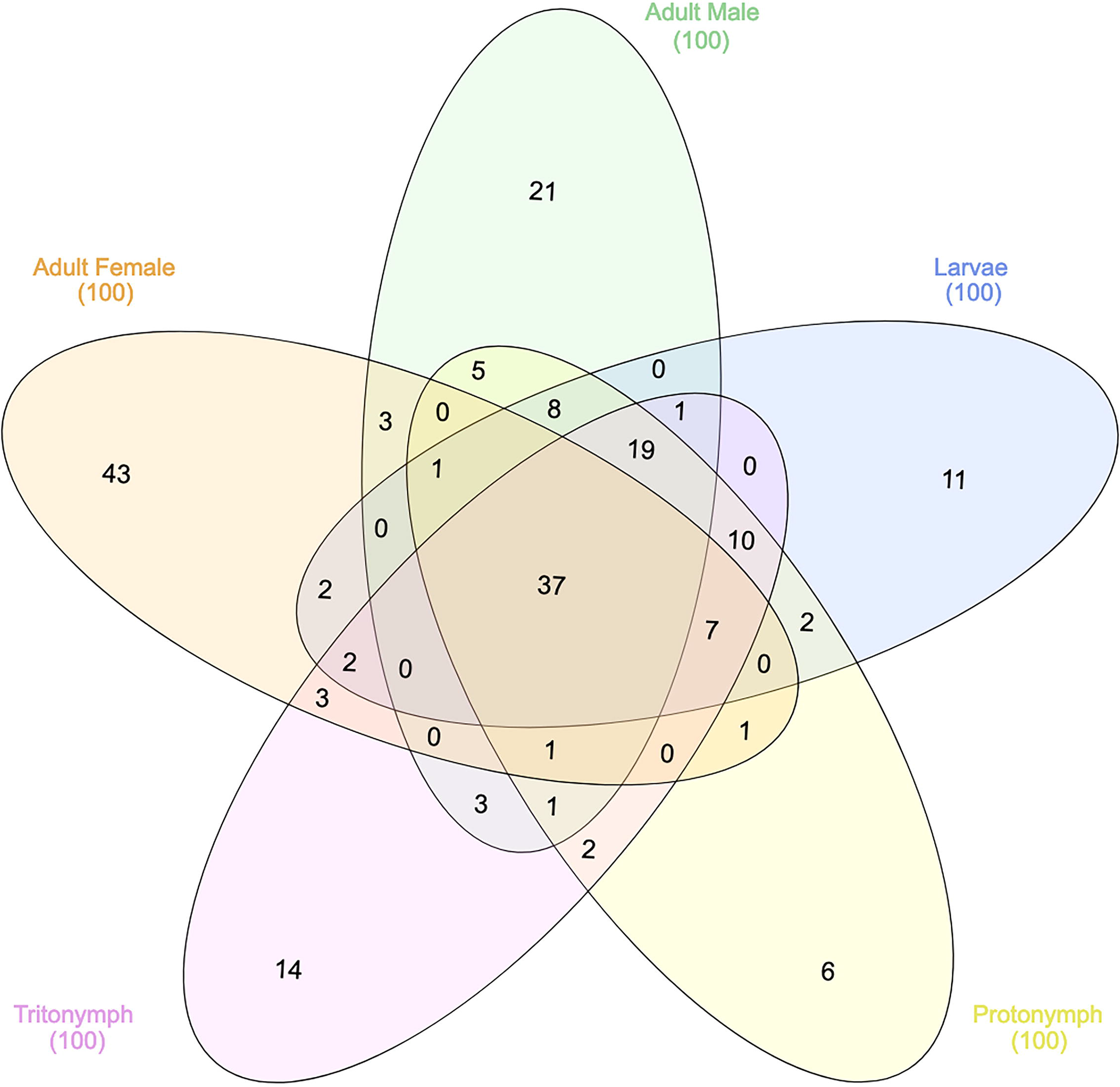
Assessment of the most abundantly expressed genes for each life-cycle stage. Five-way Venn/Euler diagram highlighting the pattern of unique and shared expression of the top 100 most abundantly expressed genes from each *P. ovis* life-cycle stage.

### Stage-specific expression of known *P. ovis* allergen genes

Homologues of a number of well-characterised allergens have been identified in *P. ovis* and many of these have been shown to elicit either pro-inflammatory, or allergic, type responses in the host and are implicated in pathogenesis of sheep scab [12,56,63]. We sought to investigate the abundance of the *P. ovis* allergens across life-cycle stages, therefore elucidating the role of each stage in the elicitation of host allergic responses. To do this we examined the expression levels of the *P. ovis* homologues of each of the 33 HDM (*D. pteronyssinus/D. farinae*) allergens currently characterised by the WHO/International Union of Immunological Societies (IUIS) across *P. ovis* life cycle stages [64,65]. *Psoroptes ovis* allergen homologues were identified for 31 of the 33 allergen classes, with just 2 classes (Der p 17 and Der p 35 (both uncharacterised allergens)) not identified in the *P. ovis* genome [66]. The relative expression of the 31 allergen homologues was analysed by hierarchical clustering and revealed unique clusters of stage-specific allergen expression (Figure 7A). Of these, one allergen (Pso o 3 (Psovi81g00190)) was expressed at very low levels (TPM ≤1) across all life-cycle stages. This gene encodes a trypsin-like serine protease and previous studies which biochemically assessed the proteolytic enzyme profiles of mixed stage *P. ovis* extracts [67] against polypeptide and peptide substrates found no evidence of serine proteinase activity [68]. The previous failure to demonstrate serine protease activity in *P. ovis* may be related to the very low levels of expression of Pso o 3, however additional serine protease-encoding transcripts were expressed in the current analysis, e.g. Pso o 6 (chymotrypsin-like serine protease) and Pso o 9 (collagenase-trypsin-like serine protease) albeit at low levels (Pso o 6: TPM ≤7, Pso o 9: TPM≤30, Figure 7A). The most highly expressed allergens included the major mite allergens Pso o 1 (a cysteine protease, mean TPM across life-cycle stages = 965), Pso o 2 (MD-2/lipid-binding protein, mean TPM = 3309), Pso o 10 (tropomyosin, mean TPM = 2658), Pso o 11 (paramyosin, mean TPM = 742), Pso o 14 (vitellogenin-apoplipophorin, mean TPM = 641), Pso o 20 (arginine kinase, mean TPM = 923), Pso o 26 (myosin light chain-like, mean TPM = 1163) and Pso o 30 (ferritin, mean TPM = 889). As shown in Figure 7A, a number of *P. ovis* allergen homologues clustered together based on their expression across specific life cycle stages. For example, Pso o 14, 27, 28, 29, 32 & 33 all showed much higher levels of gene expression in female mites than in any other stage; whilst Pso o 1, 2, 7, 8, 13, 21, 30, 34 & 36 showed highest expression in tritonymphs. Pso o 14 (vitellogenin) is a major mite allergen and the precursor for vitellin, which acts as a source of nutrients for the developing mite egg, as such elevated levels of this gene within female mites is to be expected [69–71]. As mentioned above, the putative serpin, Pso o 27, which also shows close homology to the *S. scabiei*, Sar s 27 allergen (serpin/leukocyte elastase inhibitor-like) [19,36] was highly expressed in female *P. ovis*. Although multiple copies of genes encoding Pso o 27 were identified in *P. ovis*, one transcript in particular showed significantly (p=<0.05) higher levels of expression in female mites (psovi22g04610, TPM = 15,000) than in other stages. Serpins have been implicated in arthropod development and reproduction [72] and may also influence host-pathogen interactions through immunosuppression and potential roles in innate immunity, with serpins identified in the saliva of blood-feeding ticks [73–75]. In the fruit fly, *D. melanogaster*, a Serpin-27A has been shown to inhibit the *Easter* protease, a step that is essential for the control of dorsal-ventral pattern formation in the developing embryo [76]. The near-exclusive expression of psovi22g04610 in adult female *P. ovis* mites, may suggest a potential role for this serpin-like gene in mite embryogenesis. Interestingly, the expression of the major *P. ovis* mite allergens (Pso o 1, 2, 7, 8, 13, 18, 21, 25, 30, 34 & 36) was highest in tritonymphs; these and other allergens (Pso o 1, 2, 7, 13, 14, 18, 21, 23, 27, 34 & 36) were also found to be upregulated in mixed-stage, “fed” *P. ovis* mites (Figure 7B) and may indicate that tritonymphs are the major feeding stage of *P. ovis*; potentially as part of an effort to build energy stores prior to adulthood and sexual maturation. It should be noted that protonymphs enter a stage of lethargy for up to 36 hours prior to moulting to the tritonymph stage [77] and therefore following this moult they need to acquire further nutrients before copulation, which occurs soon afterwards. More importantly the female tritonymphs moult to the adult stage just two days after the commencement of copulation and begin to lay eggs just one day later, perhaps explaining the potential increased expression of feeding related genes in the tritonymphs [77].

**Figure 7.**
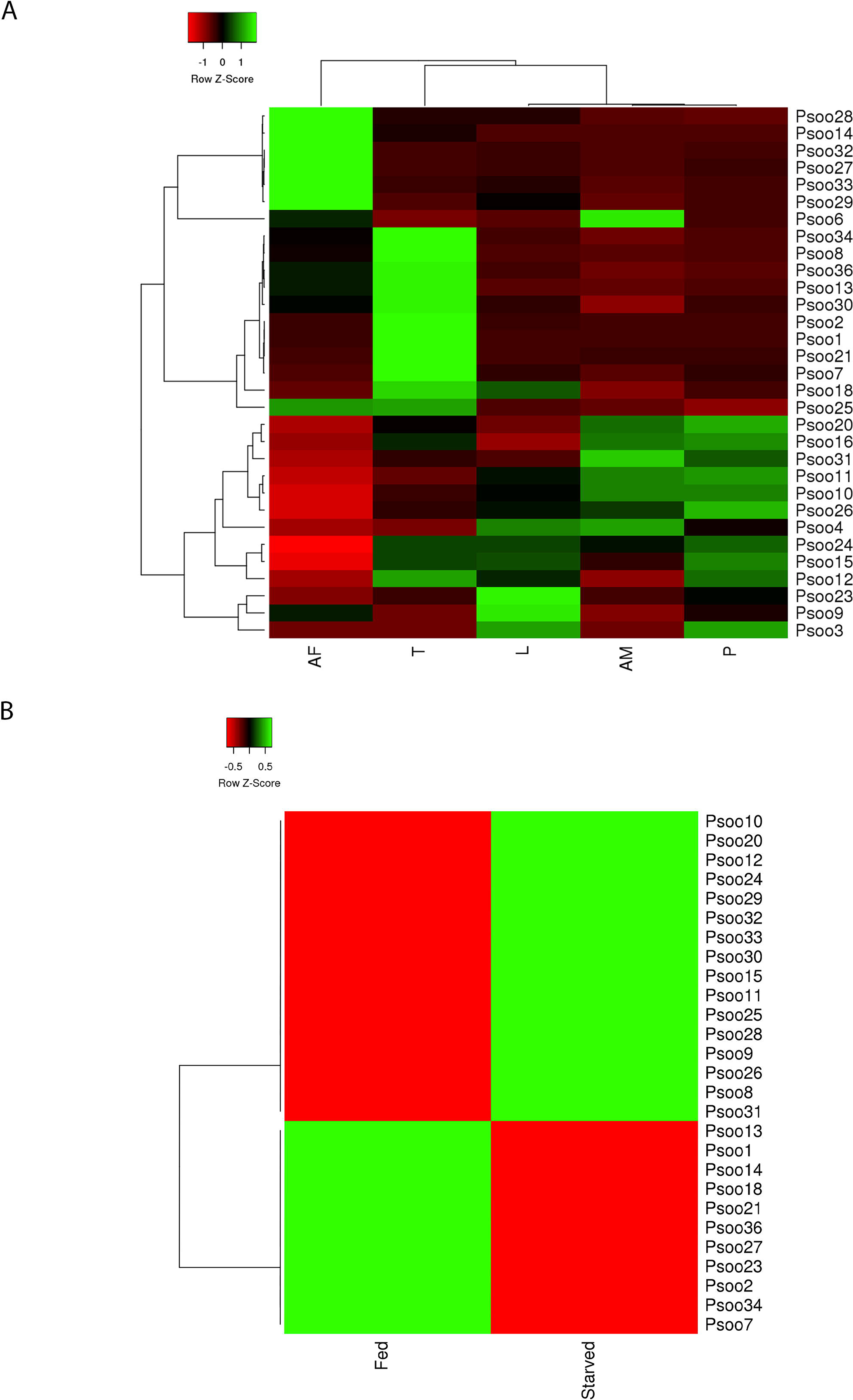
Heatmaps showing the relative expression (average TPM across three replicates/life-cycle stage/condition) of 31 characterised *P. ovis* homologous allergen genes across life-cycle stages and between “fed” and “starved” *P. ovis* mites. **A)** Stage-specific expression of *P. ovis* allergens: Larvae (L), Protonymph (P), Tritonymph (T), Adult Female (AF), Adult Male (AM). **B)** Allergen expression between “fed” and “starved” *P. ovis* mites. Each allergen is named according to the World Health Organization and International Union of Immunological Societies (WHO/IUIS) allergen nomenclature, i.e. Pso o 1 for *P. ovis* Group 1 allergen. Where multiple copies of allergen genes were identified, mean expression (TPM) data was used. Heatmaps were generated using the open source package Heatmapper [90] and Hierarchical clustering was performed using the average or unweighted pair-group method based on Pearson distance measurements.

Another cluster of *P. ovis* allergen homologues contained genes related to actin-binding and muscle contraction/motility, i.e. Pso o 10 (tropomyosin), 11 (paramyosin), 16 (gelsolin), 20 (arginine kinase), 26 (myosin light chain) & 31 (cofilin) the expression levels of which were highest in adult male mites and protonymphs. The fact that all of these genes are involved in muscle contraction and muscle motility suggests that adult males and protonymphs may be the more motile stages of *P. ovis*. This is further supported by the increased expression of Pso o 20 (arginine kinase (AK) and a homologue of the HDM allergen Der f 20 [78]) in the same cluster. AK is a phosphotransferase found in a wide variety of invertebrate species, which is especially abundant in muscle tissues, where it serves a function analogous to that of creatine kinase in vertebrates [79]. In invertebrates, AK catalyzes the reversible phosphorylation of arginine by ATP to form phosphoarginine and ADP, phosphoarginine then functions as an ATP buffer, providing high levels of ATP during periods of high cellular and/or locomotory activity [79,80]. The *P. ovis* genes attributed to each allergen class are shown in Supplementary File 3.

Our analysis identified four copies of the Group 1 cysteine protease allergen (Der p 1) homologue Pso o 1 in the *P. ovis* genome. These genes are co-located within the same scaffold and may represent a multi-gene family. The transcription data showed that three of the four copies of the gene were expressed at relatively high levels (psovi14g10400-10420 (Average TPM = 966)) whilst the remaining copy, psovi14g10430 was expressed at a very low level (TPM = 16). To further investigate this potential multi-gene family, we performed a multiple sequence alignment of the four Pso o 1 genes with the house dust mite homologues Der p 1 (*D. pteronyssinus*) and Der f 1 (*D. farinae*) (Figure 8). The alignment demonstrates the strong similarity between Der p/f 1 and Pso o 1 and also highlights the conservation of the key cysteine residues and the cysteine protease catalytic triad (QHN) as previously characterised for Der p/f 1 [81]. As detailed above Pso o 1 was most highly expressed at the tritonymph stage and was also upregulated (~4-fold) in “fed” versus “starved” *P. ovis* mites.

**Figure 8.**
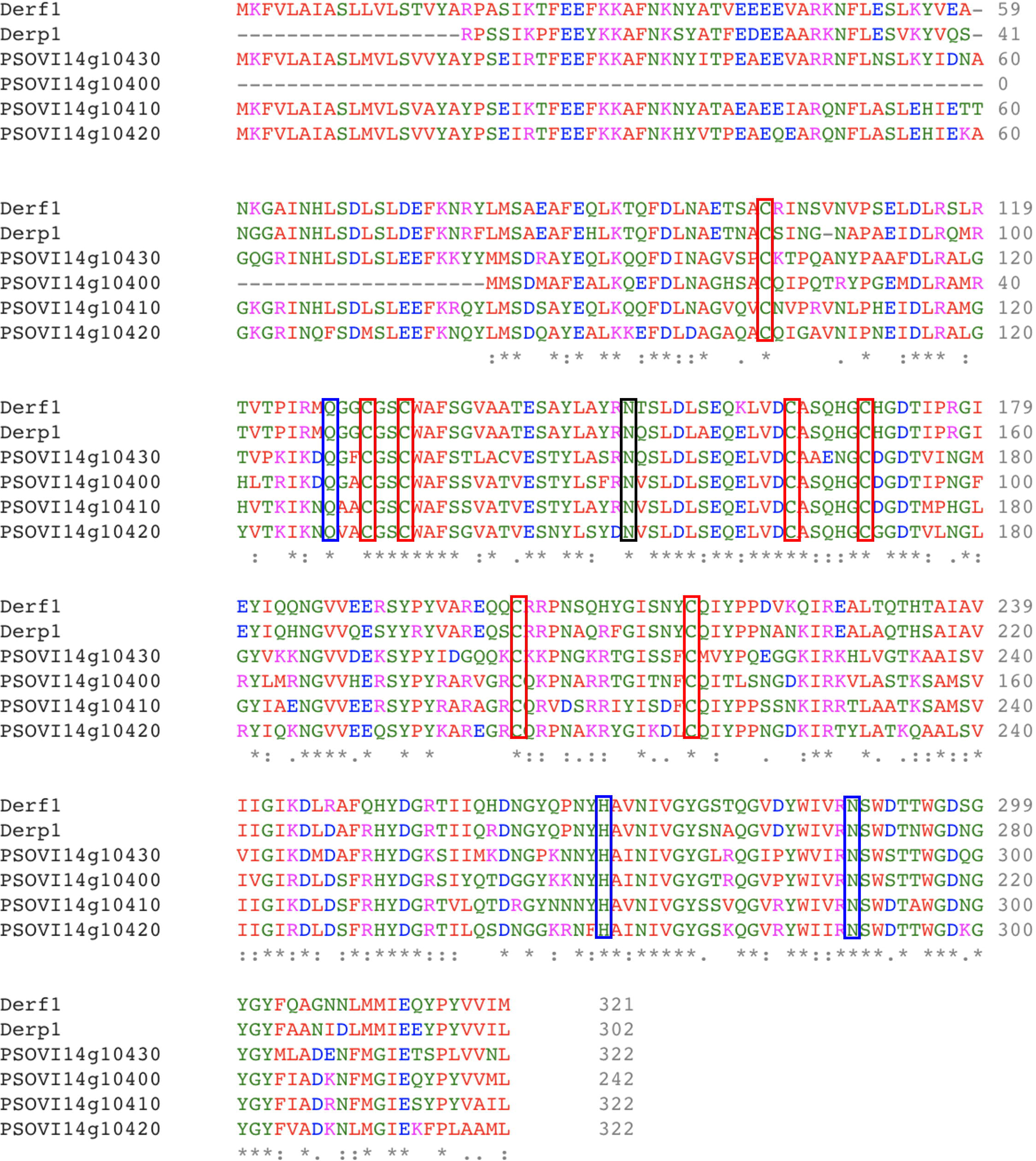
Multiple sequence alignment of Group 1 allergens from HDM (Der p 1 and Der f 1) and *P. ovis* (Pso o 1). Sequence alignment generated with EBI Clustal Omega tool [97]. Pso o 1: psovi14g10400-10430. Der f 1 (Uniprot ID: A1YW12), Der p 1 (Uniprot ID: Q3HWZ5). Red boxes mark conserved cysteine residues (x7), Blue boxes mark catalytic residues (Q, H, N) as identified in [81]. Black box marks potential Pso o 1 N-glycosylation site.

### Determination of differentially expressed genes

Pairwise analysis of differential gene expression between individual life-cycle stages and between “fed” and “starved” *P. ovis* mites was performed with edgeR (Version 3.7 [82]) based on a fold change cut-off ≥±2 and an FDR corrected p-value of ≤0.05 (Table 2). The fully annotated lists of differentially expressed genes identified from each of the pairwise life-cycle stage comparisons are available in Supplementary File 4.

**Table 2.** Numbers of differentially expressed genes (DEGs) between *P. ovis* life-cycle stages and between “fed” and “starved” mites. AF = adult females, AM = adult males, L = larvae, P = protonymph, T = tritonymph. *Direction of fold change is relative to the first stage, or condition for each comparison.

| Comparison* | Number of DEGs | Upregulated Genes | Downregulated Genes |
| --- | --- | --- | --- |
| P vs. T | 2018 | 962 | 1056 |
| P vs. L | 277 | 112 | 165 |
| L vs. T | 2734 | 1328 | 1406 |
| AM vs. T | 2908 | 1748 | 1160 |
| AM vs. P | 1590 | 1167 | 423 |
| AM vs. L | 2162 | 1451 | 711 |
| AF vs. T | 4585 | 2143 | 2442 |
| AF vs. P | 5286 | 2571 | 2715 |
| AF vs. L | 5444 | 2734 | 2710 |
| AF vs. AM | 5433 | 2376 | 3057 |
| “Fed” vs. “Starved” | 1227 | 870 | 357 |

### Feeding related gene expression in P. ovis

edgeR analysis of differential gene expression between “fed” and “starved” *P. ovis* mites identified 1,227 significantly differentially expressed genes, based on a fold change cut-off ≥±2 and an FDR-corrected p-value of ≤0.05 (Table 2). Of these the majority of genes (n=870) were upregulated in the “fed” mite population, whilst 357 genes were upregulated in the “starved” mite population (Supplementary File 4). A six-way Venn/Euler diagram analysis of gene expression across the *P. ovis* developmental stages (Larvae, Protonymph, Tritonymph, Adult Females and Adult Males with genes significantly upregulated in “fed” mites (Figure 9) demonstrated that 46% (n=242) of the tritonymph specific transcripts were also upregulated in the “fed” mite population. In contrast just 7.5% of adult male (n=67), 2.5% of adult female transcripts (n=76), 5.4% of larval transcripts (n=6) and 28% of protonymph transcripts (n=62) were found to be upregulated in “fed” mites; thus, further supporting the potential role of tritonymphs as the major feeding stage of *P. ovis*.

**Figure 9.**
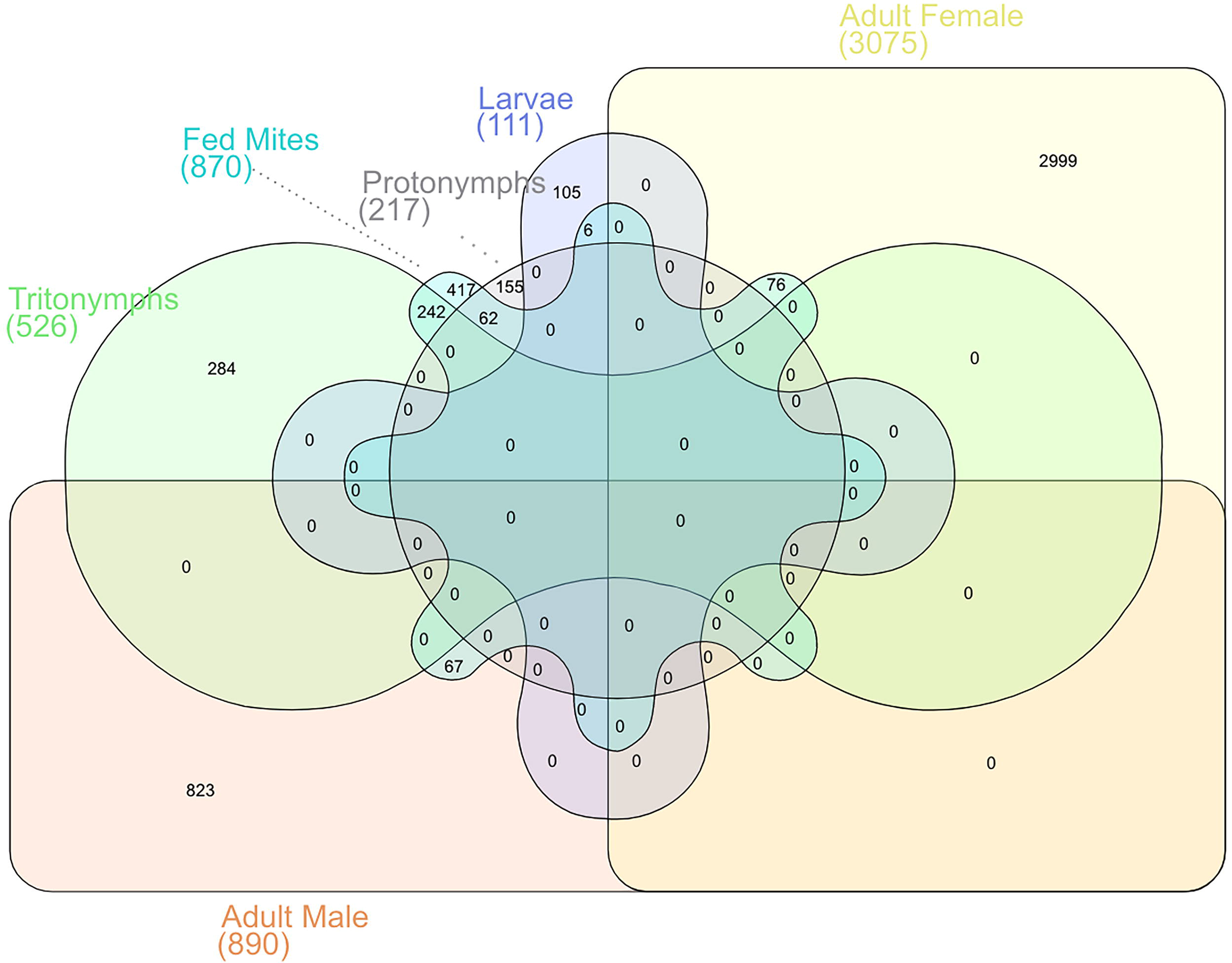
The *P. ovis* tritonymph stage is most closely associated with “fed” mite transcriptional changes. Six-way Venn/Euler diagram demonstrating gene conservation between *P. ovis* life-cycle stages. Each arm consists of the genes from the stage-specific expression clusters shown in Table 1. (*Larvae*, *Protonymph*, *Tritonymph, Adult Females and Adult Males* and genes whose expression was significantly upregulated in “fed” *P. ovis* mites (“*Fed” Mites*) compared to “starved” mites).

Amongst the 870 genes upregulated in “fed” mites, we identified 94 genes representing known allergens, including representatives of all but seven (Pso o 3, 6, 29, 31, 32, 33 and 34) of the WHO/IUIS allergen groups identified in *P. ovis*. This group included the main *P. ovis* allergens, i.e. Pso o 1, 2, 4, 7, 8, 9, 10, 11, 12, 13, 14, 15, 16, 18, 20, 21, 23, 24, 25, 26, 27, 28, 30 and 36; indicating that many allergen genes are upregulated during feeding activity, and as noted earlier, that much of this expression may be attributed to the tritonymphs. Genes representing just two allergen groups were identified in the downregulated genes, Pso o 29 (peptidyl-prolyl cis-trans isomerase or cyclophilin) and Pso o 33 (alpha-tubulin). Interestingly, our previous assessment of feeding-related expression based on a novel but limited cDNA microarray also showed down-regulation of a *P. ovis* tubulin transcript in “fed” mites [16]. A similar pattern was also observed in two previous assessments of feeding related expression in *P. ovis*: Burgess *et al.* [16] identified Pso o 1, Pso o 2, Pso o 14 and Pso o 21 as upregulated in “fed” mites, with no allergens being downregulated and McNair *et al.* [15] identified the up-regulation of Pso o 1, Pso o 27, Pso o 13 and Pso o 21 in “fed” *P. ovis*. It should be noted that both previous studies relied on the interrogation of a limited number of transcripts and were also hampered by relatively poor levels of annotation available [15,16]. Of the top upregulated genes in “fed” *P. ovis* mites, twelve belong to an uncharacterised group of senescence-associated proteins with fold change values ranging between 65 to >800-fold in “fed” mites, with the same transcripts also enriched in the tritonymph cluster (see above). As observed in the tritonymph-enriched gene cluster, the same group of five co-located genes (psovi22g03270, psovi22g03310, psovi22g03330, psovi22g03320, psovi22g03340) for which no known homologues were identified, were also found to be upregulated in the “fed” mite population with fold change values ranging from 45-92-fold higher in “fed” mites compared to “starved” mites. A number of genes encoding putative large lipid transfer proteins were also highly upregulated in “fed” mites, including vitellogenin (psovi09g01710), apolipophorin (psovi73g00070), a microsomal triglyceride transfer protein (psovi05g02390) and a homologue of the high molecular weight allergen M177 from HDMs (psovi73g00570) all of which may play a role in the transport of minerals, amino acids, lipids, and other nutrients to the developing oocyte in a range of species [83] with both vitellogenin and apolipophorin having been previously shown to be upregulated in “fed” *P. ovis* mites [16]. A number of the *P. ovis* allergens identified as being upregulated in the “fed” mites represent enzymes and may play a role in mite feeding/digestive activities, for example: Pso o 1 (cysteine protease), Pso o 4 (amylase), Pso o 9 (serine protease) and cathepsins L and B. A number of genes potentially involved in xenobiotic metabolism were also upregulated in the “fed” mite population, including seven cytochrome P450 monooxygenases (CYPs), six glutathione-S-transferases (GSTs - mu, kappa and delta-classes) and six carboxyl/choline esterases (CCEs). The upregulation of genes involved in detoxification in feeding mites is likely to be related to the increased digestive activity and subsequent higher exposure to toxic compounds from the host. We performed a Gene Ontology (GO) analysis of sequences found to be significantly upregulated in the “fed” *P. ovis* population and, of the 870 upregulated genes, significant BLAST hits and/or GO terms were assigned to 775 genes (89%) with multiple GO terms being assigned to many genes. Figure 10 shows the distribution of sequences per GO term across multiple classification levels, presented as pie-charts showing GO term distributions for Biological Process, Molecular Function and Cellular Component. GO terms for Biological Process were further subdivided between 11 categories, including: phosphate-containing compound metabolic process (18%), oxidation-reduction process (14%), response to stimulus (13%) and regulation of cellular process (9%). The Molecular Function category was further split into 12 subcategories including: hydrolase activity (19%), transferase activity (13%), oxidoreductase activity (12%), protein binding (9%) and carbohydrate derivative binding (7%). GO terms for Cellular Component were subdivided into 6 categories, including: integral component of membrane (29%), intracellular organelle part (23%) and ribosome (17%).

**Figure 10.**
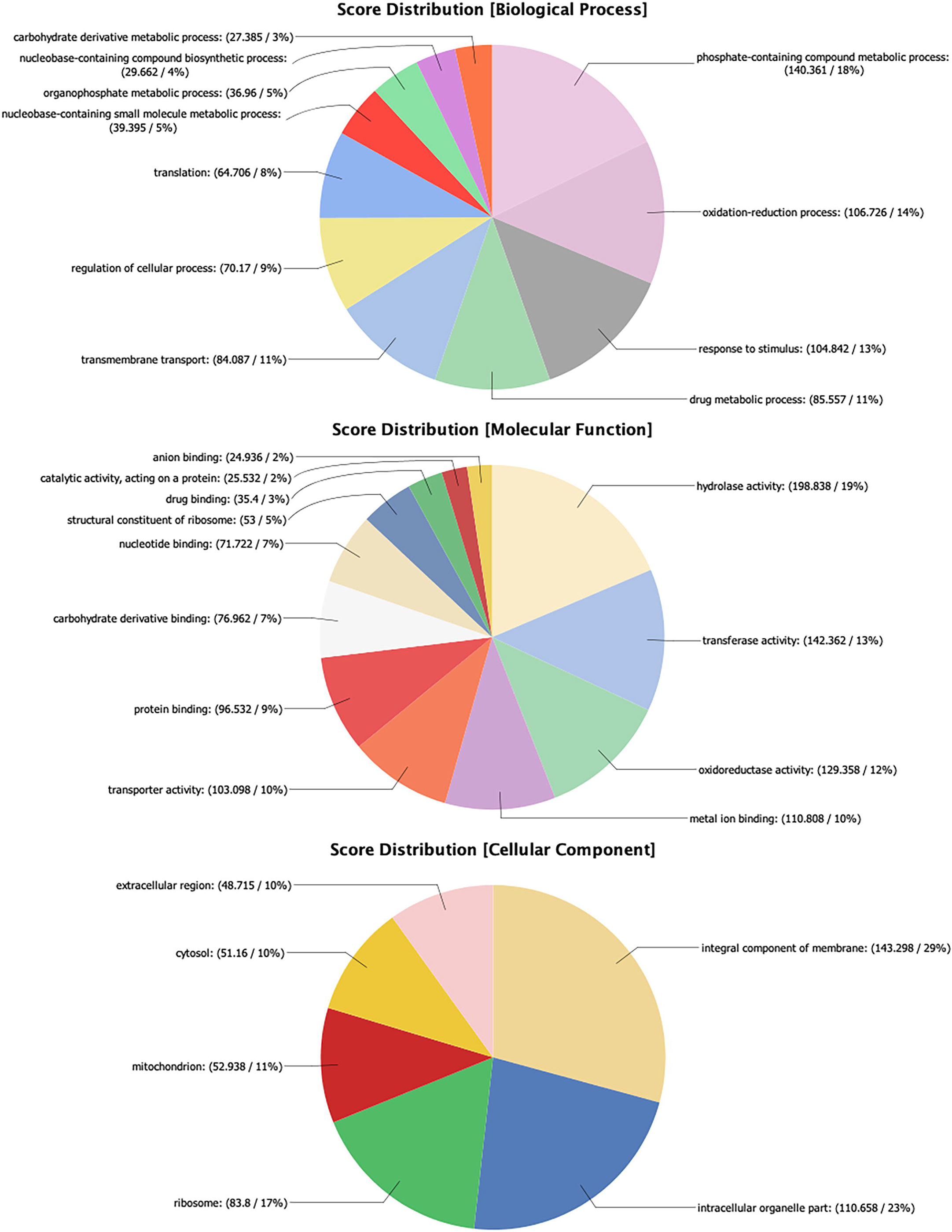
Gene ontology annotation for genes upregulated in the “fed” mite population. Each chart shows the multilevel distribution of sequences per GO term. Distribution of GO terms are summarised across three main categories: Biological Process, Molecular Function and Cellular Component.

The upregulated transcripts were further explored by mapping them to the Kyoto Encyclopedia of Genes and Genomes (KEGG) pathway database within Blast2GO; this resulted in the mapping of 213 enzyme sequences across 92 KEGG pathways. The most over-represented enzyme class was for hydrolases (42.7%) followed by oxidoreductases (22.5%), transferases (22%), lyases (6.6%), ligases (3.8%) and isomerases (2.3%). We then used Blast2GO to further investigate the enzyme code distributions between the genes upregulated in “fed” mites and those downregulated in “fed” mites. As expected more enzyme-coding genes were found in the “fed” mite population, however, these increases were most profound amongst the oxidoreductase (“starved” n=9, “fed” n=45) transferase (“starved” n=16, “fed” n=43) and hydrolase (“starved” n=23, “fed” n=61) enzyme classes. For hydrolases more sequences were found amongst the esterase (“starved” n=2, “fed” n=15) glycosylase (“starved” n=0, “fed” n=9) and peptidase families (“starved” n=2, “fed” n=15), including a cathepsin B (psovi284g00790). For transferases the main difference was observed amongst the glycosyltransferases (“starved” n=1, “fed” n=8) whilst the “fed” mites also showed an increase in the number of glutathione-S-transferases (GSTs) (“starved” n=0, “fed” n=3) a number of which are known allergens and may play a role in xenobiotic metabolism [84]. For oxidoreductases the main differences were observed in peroxidases (“starved” n=0, “fed” n=6) superoxide dismutases (“starved” n=0, “fed” n=2) indicating a potential role in response to free radicals produced by the host.

### Genes differentially expressed between P. ovis life-cycle stages

Differential expression analysis revealed a total of 7,825 genes that showed differential expression between one or more of the selected *P. ovis* life-cycle stages. We focused on the major transition phases between *P. ovis* life-cycle stages with the following comparisons being highlighted below: Adult female (AF) vs. adult male (AM), Adult female (AF) vs. tritonymph (T), Adult male (AM) vs. tritonymph (T), Larvae (L) vs. protonymph (P) and Protonymph (P) vs. tritonymph (T).

### Adult female (AF) vs. adult male (AM)

In total, 5,433 genes were found to be differentially expressed between AF and AM stages. Of these 2,376 were upregulated in the AF mites with 3,057 being downregulated with respect to the adult males. The gene with the highest degree of differential expression between AF and AM mites was a homologue of a skin secretory protein, xP2-like from *Xenopus laevis* (psovi14g01150) which showed a >900-fold increase in expression in AF mites (Average TPM = 5222 (AF) and 5 (AM)). Another gene that was highly differentially expressed was the Der p 27 homologue, Pso o 27 (psovi22g04610) with a >900-fold increase in expression in AF mites (Average TPM = 15003 (AF) and 16 (AM)). Additional female-specific genes were also highlighted in this analysis with vitellogenin (>600-fold increase in AF), vitellogenin-receptor (>87-fold increase in AF) and a *P. ovis* Group 14 allergen, apolipophorin gene (>50-fold increase in AF). A number of genes showed almost exclusive expression in the AM population compared to AFs, many of which were identified in the adult male-enriched cluster described earlier. However, significant BLAST hits were not available for most of these genes and they therefore remain as potential male-specific genes of unknown function (psovi22g06400, psovi280g06090, psovi14g05340, psovi280g02150, psovi26g00480, psovi22g04350, psovi22g04360 and psovi22g04380). Also upregulated in the AM population were the *P. ovis* allergens Pso o 10 (tropomyosin), Pso o 11 (paramyosin), Pso o 20 (arginine kinase - psovi292g02430), Pso o 26 (myosin alkali light chain), Pso o 31 (cofilin - psovi43g01240) and a *P. ovis* paxillin homologue (psovi63g00070) all of which are likely to be involved in muscle formation and contraction as described above. Finally, we identified 19 genes characterised as encoding putative secreted salivary gland peptides, of these 17 were upregulated in AM mites with just two upregulated in the AF mite population.

### Adult female (AF) vs. tritonymph (T)

The comparison between adult female and tritonymphs is important as it may inform upon differences in gene expression between the ovigerous female stage and the final nymphal stage of *P. ovis* before the nymph’s final moult into an adult. As such the tritonymph population represents the final phase of immature mites, which will go on to develop into either adult males or females. We identified 4,585 genes that were differentially expressed between AF and T stages. Of these 2,143 were upregulated in the AF mites with 2,442 being downregulated with respect to the tritonymphs. As observed earlier the AF population showed high levels of expression of a skin secretory protein xP2-like transcript with only very low levels of expression observed in the tritonymphs (635-fold higher in AF vs. T). The *P. ovis* homologue of the HDM Der f 27 allergen, a putative serpin, which was very highly expressed in the AF population was also expressed at a much lower level in the tritonymphs with a 375-fold increase in expression in the AF population compared to the tritonymphs. Other, female-related genes, such as vitellogenin (~350-fold up in AF vs. T), vitellogenin receptor (~60-fold up in AF), the Group 14 allergen, apolipophorin (Pso o 14) (147-fold up in AF) and an ABCA1 lipid exporter protein gene (72-fold up in AF) were also expressed at much higher levels in the AF population. A further group of 28 histone-related genes, including *P. ovis* homologues of histone H1B (n=2), H2A (n=2), H2B (n=3), H3, H4 (n=2), a putative histone chaperone protein, histone deacetylase (n=2), histone binding protein (n=3), a histone RNA hairpin binding protein and 12 putative histone methyltransferases were all up regulated in the AF population, indicating a potential role of chromatin remodelling during the transition phase from tritonymph to adult female. Transcripts upregulated in the tritonymphs included 42 homologues of the HDM DFP2 gene, 17 putative cuticle protein genes and 17 chitin-binding protein genes. All of these were similarly highly expressed in the tritonymph-enriched cluster described above, indicating the specific role that these genes may play in the development of the tritonymph stage. As discussed above many of the known and putative *P. ovis* allergen genes were also highly expressed in the tritonymph stage, the differential expression analysis confirmed these findings but also showed that the following antigens are often expressed at low levels in the AF mites, including: Pso o 1, Pso o 2, Pso o 7, Pso o 8, Pso o 10, Pso o 11, Pso o 13, Pso o 21, Pso o 30, Pso o 34 and Pso o 36. Two further groups of genes which showed higher levels of expression in the tritonymphs compared to the AF mites were six troponin genes, including troponin-I, C and T and the uncharacterised senescence associated proteins (n=8) identified in the tritonymph-enriched cluster above.

### Adult male (AM) vs. tritonymph (T)

As discussed above, the assessment of differential expression between adult males and tritonymphs provides an opportunity to determine gene expression changes related to the sexual maturation of the male. In total, 2,908 genes were found to be differentially expressed between AM and T stages. Of these 1,748 were upregulated in the AM mites with 1,160 being downregulated with respect to the tritonymphs. Two of the most differentially expressed genes between the AM and T stages were a putative *P. ovis* cystatin (psovi52g00910 (144-fold up in AM)) and two cathepsin-L-like genes (psovi283g01960 (12-fold up in AM) and psovi295g01310 (98-fold up in AM)). Seven further highly expressed genes (psovi22g04380, psovi280g06090, psovi280g02150, psovi14g05340, psovi22g04360, psovi22g04350 and psovi22g06400) in the AM population showed much lower expression in the T stage (ranging from 600-1500-fold higher in AM vs. T), however, these currently represent uncharacterised potentially male-enriched genes and warrant further investigation. Interestingly, a group of 13 putative testis-specific serine/threonine-protein kinases were all upregulated in the adult males compared to the tritonymphs (ranging between 2-35 fold higher in the male mites). As with the AF vs. T comparison, we observed large numbers of differentially expressed HDM, DFP2-like transcripts between the adult male and tritonymphs. However, the pattern here was different with 14 DFP2-like transcripts being upregulated in the AM mites compared to the tritonymphs, albeit all but one of these (psovi283g01710) showing relatively low levels of expression in the AM population. In contrast we identified 36 DFP2-like transcripts upregulated in the tritonymphs and many of these were expressed at very high levels (>100,000 mean TPM). Cuticle-associated protein genes (n=14) and chitin-binding proteins (n=12) were also significantly upregulated in the tritonymphs compared to the adult males, with just one cuticle protein and three chitin-binding factors up regulated in the AM population. In terms of allergen expression, the majority of allergens were again upregulated in the tritonymphs compared to the adult males, however, there were a few exceptions, with Pso o 10 (tropomyosin), Pso o 11 (paramyosin), Pso o 20 (arginine kinase), Pso o 26 (myosin alkali light chain) & Pso o 31 (cofilin) all upregulated in the AM population and all with potential roles in muscle development and contraction.

### Larvae (L) vs. protonymph (P)

We identified 277 genes that were differentially expressed between L and P stages. Of these 165 were upregulated in the protonymphs with 112 being downregulated with respect to the larvae. Of the 165 upregulated genes in the protonymphs, 29 of these (18%) showed homology to HDM, DFP2-like factors, many of which showed very high levels of expression in the protonymphs with low levels of expression in the larvae. In addition, cuticle-associated factors (n=15) were also significantly upregulated in the protonymphs with just one cuticle-like protein upregulated in the larvae. This seems to indicate that the *P. ovis* DFP2-like genes and the high levels of cuticle-like protein gene expression are more-associated with the nymphal stages, rather than the larval stage.

### Protonymph (P) vs. tritonymph (T)

In total, 2,018 genes were found to be differentially expressed between P and T stages. Of these 962 were upregulated in the protonymphs with 1,056 being downregulated with respect to the tritonymphs. Of these 44 differentially expressed transcripts relate to HDM, DFP2-like genes, with 21 of these being upregulated in the tritonymphs and 23 upregulated in the protonymphs; suggesting that there may be different families of DFP2-like genes, which share similar stage-specific patterns of expression. To investigate this further we generated an alignment of the predicted protein sequences for the 44 differentially expressed DFP2-like genes and also for all of the predicted *P. ovis* DFP2-like proteins identified in the genome sequence (n=81). The resulting phylogenetic trees (presented as circular phylograms) can be seen in Figure 11. The first of these (Figure 11A) shows the differentially expressed (protonymph vs. tritonymph) DFP2-like protein sequences, whilst Figure 11B shows all 81 predicted DFP2-like protein sequences identified in the *P. ovis* genome. The circular phylograms clearly demonstrate clustering of the DFP2-like sequences that are specifically upregulated in the tritonymphs (labelled with a red T) and, separately, those upregulated in protonymphs (labelled with blue P), demonstrating conserved sequences in particular life-cycle stages and suggesting the potential for distinct biological functions of DFP2-like proteins in each stage. In addition, of the 21 DFP2-like genes upregulated in tritonymphs, 16 are co-located on the same scaffold (psovi283) indicating the potential for these to be co-regulated or expressed. Similar patterns were also observed for cuticle protein genes (n=9) six of which were upregulated in protonymphs, with three upregulated in tritonymphs, two of which were co-located on a single scaffold (psovi22g00570 and psovi22g00580). Chitinases and chitin-binding proteins (n=7) also showed a mixed pattern of expression, with four upregulated in tritonymphs and 3 upregulated in protonymphs. Homeobox factors were mostly upregulated in the protonymph stage with 17 related transcripts, of which 15 were upregulated in the protonymphs, including two putative *P. ovis* developing brain homeobox protein 1 (DBX1) homologues, co-located on a single scaffold (psovi43g01050 and psovi43g01060), which may play a role in central nervous system patterning in *D. melanogaster* [85]. A large number of putative allergen genes showed increased expression in the tritonymph stage, including, Pso o 1 (all four copies), Pso o 2, Pso o 7, Pso o 8, Pso o 14 and Pso o 36, whilst Pso o 10 and Pso o 11 were upregulated in the protonymphs. Also upregulated in the tritonymphs were three putative *P. ovis* cystatin genes, six *P. ovis* cathepsins (B and L) and 12 copies of the novel *P. ovis* senescence associated proteins.

**Figure 11.**
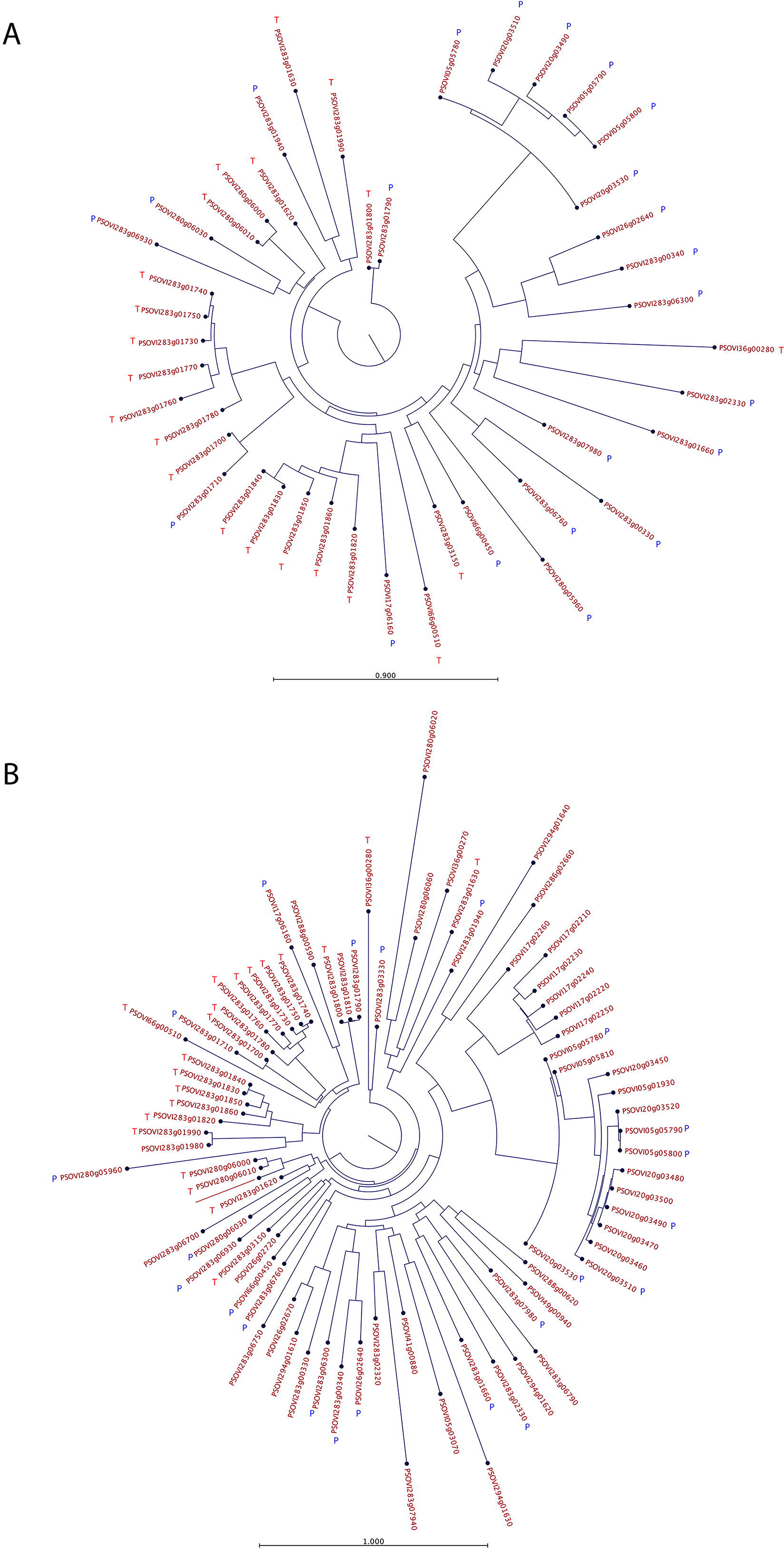
Phylogenetic analysis of DFP2-like proteins from *P. ovis*. Trees were constructed using a maximum likelihood phylogeny method with neighbour joining tree construction and Jukes-Cantor distance measure with 100 bootstraps. **A)** DFP2-like genes differentially expressed (n=44) between protonymph and tritonymph stages. **B)** The 81 predicted *P. ovis* DFP2-like protein sequences from the draft *P. ovis* genome. DFP2-like genes upregulated in tritonymphs (vs. protonymphs) are highlighted with a red T; whilst those upregulated in protonymphs (vs. tritonymphs) are highlighted with a blue P.

## Conclusions

This study represents the first large-scale genomic and transcriptomic analysis of sex- and stage-specific and feeding-related gene expression in a parasitic astigmatid mite. It also presents, for the first time the complete annotation of the *P. ovis* draft genome and predicted transcriptome. The analysis showed clear patterns of gene expression attributed to individual *P. ovis* life cycle stages including, for the first time, the demonstration that previously-characterised allergens may exhibit both stage-specific and feeding-related patterns of gene expression; a finding of particular importance when developing novel means of control, i.e. vaccination. Stage-specific allergen expression was demonstrated for a number of known *P. ovis* allergens, including the major mite allergen Pso o 1 (a homologue of the HDM allergen Der p 1). By exploiting the draft genome for *P. ovis* we were also able to show that Pso o 1 is a member of a multigene family (with four copies co-located on a single scaffold) which shows high levels of expression across life-cycle stages but with increased expression in tritonymphs. The observation that tritonymph stage-enriched expression overlaps closely with feeding-related gene expression points to the tritonymphs being the main feeding stage of *P. ovis* and the identification of novel multigene families (i.e. *P. ovis* putative senescence-associated proteins) may offer future targets for control. The analysis of sex-specific expression in *P. ovis* showed that large numbers of transcripts expressed in the female mites are dedicated to the process of oogenesis, whilst in the males we observed expression of numerous genes involved in muscle development and contraction, perhaps reflecting the increased mobility of the adult males as they seek out multiple females for copulation. We identified a further novel multigene family in *P. ovis*, which consisted of 81 genes showing close homology to the HDM DFP2-like gene. Phylogenetic analysis of *P. ovis* DFP2-like genes revealed separate clustering of these DFP2-like sequences that were specifically upregulated in protonymphs and tritonymphs, demonstrating expression of closely-related members of this family in particular *P. ovis* stages and suggesting distinct biological function amongst the DFP2-like proteins. By building on our recent publication of the *P. ovis* draft genome we have now generated the first genomic and transcriptomic atlas of gene expression in *P. ovis*. This represents a unique resource for this economically important parasite and also provides the first gene expression atlas for an astigmatid parasitic mite, which can now be exploited by the wider acarid-research community. The OrcAE platform and accompanying *P. ovis* transcriptomic atlas, is publicly accessible and represents a means by which the draft *P. ovis* genome can be further improved via a process of community-led manual curation.

## Methods

### P. ovis collection and life-cycle staging

Ethical approval for the study was obtained from the Moredun Research Institute Experiments Committee [E03/17]. *Psoroptes ovis* mites (a mixed population consisting of adults, nymphs and larval stages) were harvested from infested donor sheep maintained at the Moredun Research Institute as previously described [18]. Individual *P. ovis* mites derived from this mixed population were separated into the following life-cycle stages by staff at Fera Science Ltd: Adult Females (AF), Adult Males (AM), Larvae (L), Protonymph (P) and Tritonymph (T). Mites from each life-cycle stage were then divided into three equal sized pools, snap frozen in liquid nitrogen and stored at −80°C prior to RNA extraction.

### P. ovis collection – “fed” and “starved” mites

The “fed” *P. ovis* mite samples (n=3) were taken from the mixed population as described above, prior to staging and split into 3 pools. “Starved” *P. ovis* mites (n=3) were also obtained from the mixed population but following the harvest, mites (~100mg) were placed into a 75cm^2^ vented cap cell culture flask (Corning, UK) and incubated for 4 days at 25°C with 80-90% relative humidity and then split into 3 pools.

### RNA extraction and quality control

Total RNA was extracted from the triplicate pools for each life-cycle stage and from the “fed” and “starved” mite populations. This was achieved by homogenisation (within a pestle and mortar under liquid nitrogen) in TRIzol Reagent (Thermo Fisher Scientific Ltd, UK) according to the manufacturer’s protocol. RNA samples were further purified using a Qiagen RNeasy kit, following the manufacturer's RNA clean-up protocol and on-column DNase I digestion for 15 minutes at room temperature, prior to elution into RNase free dH2O. Total RNA yield was assessed on a Nanodrop spectrophotometer (Nanodrop, Thermo Scientific Ltd, UK) and RNA quality was determined on an Agilent Bioanalyser (Agilent, UK) using the RNA Nano-chip kit (Agilent, UK).

### Library preparation and transcriptome sequencing

TruSeq RNA-seq libraries (Illumina, San Diego, USA) were prepared from the 15 *P. ovis* life-cycle stage RNA samples (in triplicate for each of the life-cycle stages (biological replicates)) and for the six *P. ovis* samples for the “fed” vs. “starved” comparison (3 pools of each) according to the manufacturer’s instructions. Sequencing was performed on the Illumina HiSeq 2000 platform with version 3 chemistry (Ilumina, USA) by the Gene Pool Next Generation Sequencing and Bioinformatics Service at the University of Edinburgh with 50 base single-end sequencing for the *P. ovis* life-cycle stages and 50 base paired-end sequencing for the “fed” vs “starved” mite comparison.

### Bioinformatic analysis

Base calls were made using the Illumina CASAVA 1.8 pipeline. Post-sequencing, read quality of raw FASTQ files was checked with FastQC v0.10.154. The CLC Genomics Workbench (Version 12, Qiagen Ltd) was used for adapter, quality, ambiguity, and length trimming. For alignment of the read data, we employed the draft genome assembly for *P. ovis* which is a ∼63.2-Mb genome containing 12,041 predicted protein-coding genes [15]. Pseudo alignment of the read data to the *P. ovis* genome (Accession ID: PQWQ01000000)[18] was performed in Kallisto (Version 0.44.0 [26]). Kallisto is a novel transcriptome-based quantification package that avoids the considerable bias that can be introduced by a genome alignment step [86]. Kallisto generated read counts (transcripts per million (TPM)) for all RNA-seq samples, which were used as input for the network clustering within the Graphia Professional package (Version 2.0, Kajeka, Edinburgh, UK, formerly BioLayout Express 3D [29,87]). Functional annotation of the *P. ovis* genome and specific-gene clusters was performed within the Blast2GO package (Version 5) [88]. Venn/Euler diagram analysis was performed using InteractiVenn [89]. Heatmaps were generated using the open source package Heatmapper [90].

### Determination of differentially expressed genes

The statistical package edgeR (Version 3.7) within the R software suite (Version 3.1) was used to analyse the RNA-seq (Illumina Hi-Seq) data and to identify transcripts significantly differentially expressed between *P. ovis* life-cycle stages and between “fed” and “starved” mites [82,91]. Read count data, as TPM from the Kallisto package, for each replicate of each stage, or condition, were used as the input data for the differential expression analysis. By default, edgeR uses the number of mapped reads (in this case TPM column sums) and estimates a normalisation factor taking into account sample-specific effects, these factors are then combined and used as an offset in the negative binomial model within edgeR [92]. As a pre-filtering step we also removed features without a TPM value of >1 in each of the 3 replicates per life-cycle stage or condition. Significantly differentially expressed transcripts were classified as those having a fold change ≥±2.0 between each of the pairwise comparisons of *P. ovis* life-cycle stages (n=10) or between “fed” and “starved” mites and a False Discovery Rate (FDR) corrected p-value of ≤0.05 [93]. Putative functions were assigned to the differentially expressed transcripts following homology searches using the NCBI, Basic Local Alignment Search Tool (BLAST) against the NCBI non-redundant (nr) database and motif identification using IPS within the Blast2GO package (Version 5) [88].

### Phylogenetic assessment of the predicted protein sequences of the *P. ovis* DFP2-like genes

Multiple sequence alignments were generated using a progressive alignment algorithm within the CLC Genomics Workbench (Version 12, Qiagen Ltd) [94]. The multiple sequence alignments were then used to produce phylogenetic trees based on a maximum likelihood phylogeny method with neighbour joining tree construction and Jukes-Cantor substitution model with 100 bootstraps, within the CLC Genomics Workbench (Version 12, Qiagen Ltd).

### Interactive web-based presentation of the *P. ovis* genome and gene expression atlas

*Ab initio* gene predictions and annotation of the *P. ovis* genome were performed using the gene prediction platform, EuGene, as previously described [18]. Predicted genes were functionally annotated through a combination of InterProScan and reciprocal best BLAST hits. Predicted secretion signals, transmembrane helices, and other functional domains were generated within PHOBIUS [95,96]. The full annotation for each *P. ovis* gene has now been made available within the Online Resource for Community Annotation of Eukaryotes (OrcAE) along with gene expression (RNA-seq) data from the current study viewable as a gene expression atlas.

## Accession numbers

The sequence data produced and analysed in this study is fully compliant with the MINISEQE guidelines and was deposited in the publicly accessible NCBI Sequence Read Archive (SRA) Database under the project accession number PRJNA521406. The *P. ovis* draft genome sequence is available at DDBJ/ENA/GenBank under the accession number PQWQ01000000. The full annotation of the *P. ovis* genome has been made publicly available via the Online Resource for Community Annotation of Eukaryotes (OrcAE) via the following link: https://bioinformatics.psb.ugent.be/orcae/

## Acknowledgments

This work was supported by the UK Department for Environment Food and Rural Affairs (Defra) under research projects OD0555 & OD0556 and also received financial support from the Scottish Government’s Rural Affairs Food and Environment Strategic Research (RESAS) programme. The Moredun Research Institute is one of the Scottish Government’s major research providers under the collective of the Scottish Environment, Food and Agriculture Research Institutes (SEFARI). The authors would like to thank the Gene Pool Next Generation Sequencing and Bioinformatics Service at the University of Edinburgh for the *P. ovis* mite transcriptome sequencing. We also acknowledge the excellent service and animal care provided by the Moredun Research Institute, Bioservices team.

---

**Table S1.** Total number of Illumina Solexa Hi-Seq reads for each of the fifteen RNA samples from *P. ovis* life-cycle stages and for “fed” (F) and “starved” (S) mites. AF = adult females, AM = adult males, L = larvae, P = protonymph, T = tritonymph.

| Sample Description | Sample ID | Total Reads |
| --- | --- | --- |
| Adult Females | AF_1 | 12,660,138 |
| Adult Females | AF_2 | 16,172,943 |
| Adult Females | AF_3 | 15,960,093 |
| Adult Males | AM_1 | 13,822,039 |
| Adult Males | AM_2 | 17,998,180 |
| Adult Males | AM_3 | 9,886,599 |
| Larvae | L_1 | 14,670,039 |
| Larvae | L_2 | 12,066,561 |
| Larvae | L_3 | 13,094,220 |
| Protonymphs | P_1 | 11,038,552 |
| Protonymphs | P_2 | 19,996,884 |
| Protonymphs | P_3 | 13,850,774 |
| Tritonymphs | T_1 | 26,140,689 |
| Tritonymphs | T_2 | 10,155,712 |
| Tritonymphs | T_3 | 8,524,992 |
| “Fed” | F_1 | 9,454,291 |
| “Fed” | F_2 | 16,091,427 |
| “Fed” | F_3 | 12,419,967 |
| “Starved” | S_1 | 12,851,430 |
| “Starved” | S_2 | 13,229,126 |
| “Starved” | S_3 | 8,183,501 |

